# Single-Molecule FRET Illuminates Structural Subpopulations and Dissects Crucial Molecular Events During Phase Separation of a Prion-Like Low-Complexity Domain

**DOI:** 10.1101/2023.05.23.541917

**Authors:** Ashish Joshi, Anuja Walimbe, Anamika Avni, Sandeep K. Rai, Lisha Arora, Snehasis Sarkar, Samrat Mukhopadhyay

## Abstract

Biomolecular condensates formed via phase separation of proteins and nucleic acids are thought to be associated with a wide range of cellular functions and dysfunctions. We dissect critical molecular events associated with phase separation of an intrinsically disordered prion-like low-complexity domain of Fused in Sarcoma by performing single-molecule studies that permit us to access the wealth of molecular information that is skewed in conventional ensemble experiments. Our single-molecule FRET experiments reveal the coexistence of two conformationally distinct subpopulations in the monomeric form. Single-droplet single-molecule FRET studies coupled with fluorescence correlation spectroscopy, picosecond time-resolved fluorescence anisotropy, and vibrational Raman spectroscopy indicate that structural unwinding switches intramolecular interactions into intermolecular contacts allowing the formation of a dynamic network within condensates. A disease-related mutation introduces enhanced structural plasticity engendering greater interchain interactions that can accelerate pathological aggregation. Our findings provide key mechanistic underpinnings of sequence-encoded dynamically-controlled structural unzipping resulting in biological phase separation.

## Introduction

Living cells compartmentalize their biochemical components and processes using well-defined membrane-bounded organelles. A growing body of rapidly evolving research reveals that in addition to such conventional membrane-bounded organelles, cells contain noncanonical membraneless organelles that are thought to be formed via liquid-liquid phase separation of proteins, nucleic acids, and other biomolecules^1–10^. These membraneless compartments, also termed biomolecular condensates, include nucleolus, stress granules, P-bodies, Cajal bodies, nuclear speckles, and so on. These membraneless bodies are highly dynamic, liquid-like, regulatable, permeable, nonstoichiometric supramolecular assemblies involved in the spatiotemporal regulation of vital cellular processes including genome organization, RNA processing, signaling, transcription, stress regulation, immune response, and so forth^11–13^. Recent studies have established that intrinsically disordered proteins/regions (IDPs/IDRs) possessing low-complexity and prion-like domains are the key candidates for biological phase separation^7, 8, 14–16^. These studies revealed that the presence of low-sequence complexity promotes intrinsic disorder, conformational flexibility, structural heterogeneity, and multivalency that enable the polypeptide chains to participate in a multitude of ephemeral chain-chain interactions governing the making and breaking of noncovalent interactions on a characteristic timescale. These noncovalent intermolecular interactions involve electrostatic, hydrophobic, hydrogen bonding, dipole-dipole, π–π, and cation–π interactions and yield a highly dynamic liquid-like behavior of phase-separated biomolecular condensates^17–21^. Biomolecular condensate formation involves a density transition coupled to the percolation that results in the dense phase comprising a viscoelastic network fluid^4^. Such condensates comprising viscoelastic fluids can undergo aberrant liquid-to-solid phase transitions resulting in the maturation and hardening of these assemblies into solid-like aggregates that are associated with a wide range of neurodegenerative diseases^2, 12–14, 22^.

An archetypal phase-separating protein, Fused in Sarcoma (FUS), is a highly abundant protein belonging to the FUS family of proteins. Liquid-like condensates of FUS are thought to play crucial roles in RNA processing, DNA damage repair, paraspeckle formation, miRNA biogenesis, and the formation of stress granules. On the contrary, solid-like aggregates of FUS are identified as pathological hallmarks of several neurodegenerative diseases including Amyotrophic Lateral Sclerosis (ALS) and Frontotemporal Dementia (FTD)^23–27^. FUS exhibits a multidomain architecture comprising an intrinsically disordered N-terminal domain and a partly structured C-terminal RNA binding domain. The N-terminal domain contains a QSGY-rich prion-like low-complexity domain, whereas, the C-terminal RNA-binding domain (FUS-RBD) consists of an RNA-recognition motif (RRM), two RGG-rich stretches, a zinc finger domain, and a short nuclear localization signal (NLS) (Fig. 1a). The N-terminal, intrinsically disordered, prion-like low-complexity domain, termed FUS-LC, has been identified as the major driver of self-assembly into liquid-like condensates, hydrogels, and solid-like aggregates (Fig. 1b)^24–28^. Previous studies have established that FUS-LC serves as a model prion-like system to investigate the fundamental biophysical principles of phase separation and maturation. Structural characterizations have indicated that FUS-LC remains intrinsically disordered in both monomeric dispersed and condensed phases^24, 29–35^. However, the complex interplay of the key molecular determinants and the crucial molecular events that critically govern the course of macromolecular phase separation of FUS-LC remain elusive. The key question of how sequence-encoded conformational plasticity, structural distributions, and chain dynamics control weak, multivalent, transient intermolecular interactions resulting in the formation of liquid-like condensates is of paramount importance both in normal cell physiology and disease biology.

**Fig. 1.**
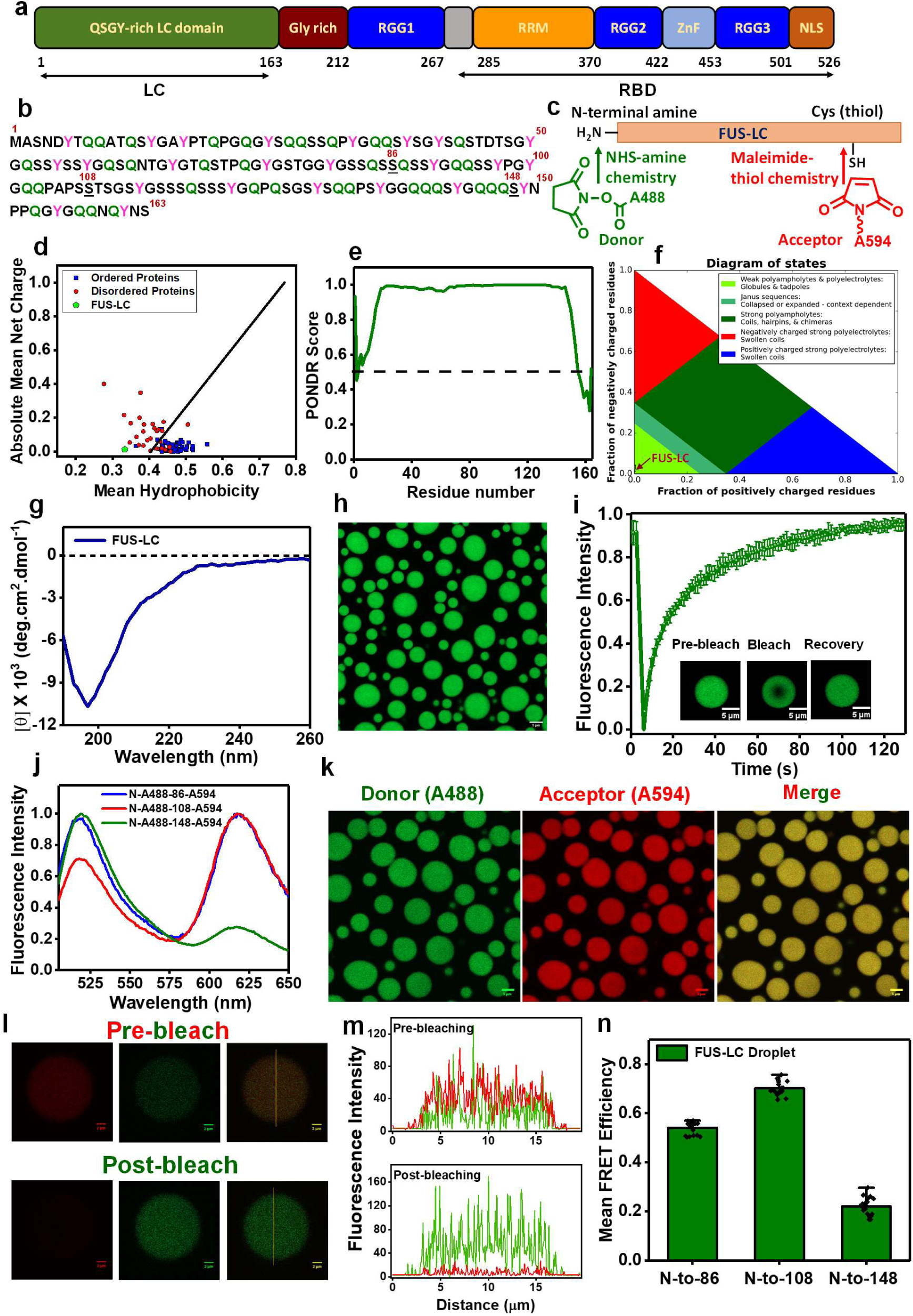
Sequence architecture and characterization of FUS-LC liquid condensates. a. Domain architecture of full-length FUS. b. The amino acid sequence of FUS-LC highlighting the tyrosine and glutamine residues. Residue positions for single-cysteine mutants are underlined. c. Schematic of orthogonal labeling chemistry utilized for site-specific labeling of N-terminal with NHS ester of AlexaFluor488 and cysteine with thiol-active AlexaFluor594-maleimide. d. The mean hydrophobicity vs. the mean net charge for a range of natively ordered and disordered proteins, FUS-LC is represented in green. e. Predictor of Natural Disordered Regions (PONDR) showing unstructured FUS-LC. f. A sequence annotated diagram of IDP states based on the fraction of negative and positive charged residues shows FUS-LC as compact globules and tadpole-like conformations g. A far-UV CD spectrum of FUS-LC indicating random-coil conformation. h. Confocal image of FUS-LC droplets containing 0.1 % AlexaFluor488-labeled protein. i. FRAP kinetics of multiple droplets (n = 12) represented by the mean and standard deviation (0.1% AlexaFluor488-labeled FUS-LC was used for FRAP measurements). j. Ensemble steady-state fluorescence emission spectra of dual-labeled FUS-LC showing donor and acceptor fluorescence upon donor excitation at 488 nm. k. Two-color Airy scan confocal image of FUS-LC droplets formed in the presence of 0.05% FUS-LC labeled with AlexaFluor488 at the N-terminal and AlexaFluor594 at Cys 86 (N-A488-86-A594) l. A representative image of acceptor photobleaching FRET of individual dual-labeled FUS-LC condensates and fluorescence intensity profiles (m) showing a decrease in the acceptor fluorescence intensity and an increase in the donor fluorescence intensity upon acceptor photobleaching. n. Mean FRET efficiencies in FUS-LC condensates were estimated from the acceptor photobleaching (n = 19 droplets) for three constructs varying the intramolecular distance, namely, N-to-86, N-to-108, and N-to-148.

In this work, we elucidate the structural heterogeneity, distributions, and interconversion dynamics of FUS-LC using single-molecule experiments that permit us to monitor the conformational states in a molecule-by-molecule manner. Such single-molecule experiments offer a powerful approach to accessing the incredible wealth of molecular information that is normally skewed in conventional ensemble-averaged experiments^36–42^. These single-molecule studies allow us to interrogate one molecule at a time and directly capture the hidden conformational states, characteristic conformational fluctuations, and interconversion dynamics. We utilized highly sensitive single-molecule FRET (Förster resonance energy transfer) using FUS-LC constructs site-specifically and orthogonally labeled with a donor-acceptor pair (Fig. 1c) that allowed us to detect and characterize structurally distinct states and conformational distributions within structurally heterogeneous populations in the monomeric dispersed phase and the protein-rich condensed phase. Our single-molecule FRET studies varying the inter-residue distance in conjunction with fluorescence correlation spectroscopy (FCS), picosecond time-resolved fluorescence anisotropy, and vibrational Raman spectroscopy within individual phase-separated condensates permitted us to dissect the conformational shapeshifting events associated with phase separation of FUS-LC. We also elucidated the impact of a clinically-relevant pathological mutation on the conformational distribution and dynamics that alters the phase behavior of the low-complexity domain.

## Results

### Single-droplet FRET imaging hints at long-range intramolecular interactions in the condensed phase

FUS-LC comprising 163 residues exhibits a near-uniform distribution of amino acids, serine (S), tyrosine (Y), glycine (G), and glutamine (Q), and is characterized by a low net charge and low mean hydrophobicity (Fig. 1b, d). We began with the bioinformatics characterization using PONDR^43^, which confirmed the presence of intrinsic disorder in FUS-LC (Fig. 1e). IDPs often possess a sequence composition that is characterized by a low mean hydrophobicity and a high net charge^44^. However, FUS-LC carries a low net charge with an NCPR value < 0.25; such a polypeptide can exist as compact globular ensembles in contrast to the well-solvated expanded conformations exhibited by charged IDPs^45, 46^. Based on the charge composition, FUS-LC is predicted to adopt a compact^47^ or tadpole-like^45^ structure, as also evident from the diagram of states, which predicts IDP conformations based on the fraction of positively and negatively charged residues within the sequence (Fig. 1f). To experimentally validate, we recombinantly expressed and purified FUS-LC and performed circular dichroism (CD) spectroscopic measurements of monomeric FUS-LC which exhibited a characteristic disordered state (Fig. 1g). Upon addition of salt, as observed previously^29–31^, a homogeneous solution of FUS-LC underwent phase separation as indicated by a rise in the turbidity of the solution (Supplementary Fig. 1a). Next, in order to directly visualize the droplet formation, we took advantage of the fact that FUS-LC does not contain any lysine residue and selectively labeled the N-terminal amine using the amine-reactive AlexaFluor488 succinimidyl ester (NHS ester) (Fig. 1c). This labeling strategy also allowed us to perform (selective) orthogonal dual labeling for FRET studies (see below). Using AlexaFluor488-NHS-labeled FUS-LC, we imaged the droplets using confocal microscopy (Fig. 1h). These droplets exhibited liquid-like behavior as evident by rapid and complete fluorescence recovery after photobleaching (FRAP) (Fig. 1i). These observations indicated that FUS-LC undergoes liquid phase condensation upon the addition of salt and are in agreement with previous reports^24, 29, 30^. Next, in order to elucidate the conformational changes associated with phase separation, we performed intramolecular FRET measurements both in monomeric dispersed and condensed phases.

The FRET donor (AlexaFluor488) was installed using the N-terminal NHS chemistry and the acceptor (as stated above), whereas, thiol-active AlexaFluor594-maleimide, was covalently linked at a Cys position of the single-Cys variants created along the FUS-LC polypeptide chain (Fig. 1b, c). We created three single-Cys mutants of FUS-LC (Cys residues at residue positions 86, 108, and 148) that encompassed the significant part of the polypeptide chain from the N-to the C-terminus and allowed us to record three intramolecular distances from the N-terminal end (N-to-86, N-to108, and N-to-148). The orthogonal labeling chemistries (NHS labeling at N-terminal amine and thiol-maleimide chemistry at Cys residues) yielded three FRET constructs selectively labeled with a donor (AlexaFluor488) and an acceptor (AlexaFluor594) (Fig. 1c). As a prelude to performing more advanced single-molecule FRET experiments, we carried our ensemble steady-state FRET measurements both in spectroscopy and microscopy formats. The dual-labeled FUS-LC constructs exhibited energy transfer both in the monomeric dispersed state (Fig. 1j) and in the droplet phase as evident by an overlapping two-color confocal microscopy image (Fig. 1k). We next performed single-droplet acceptor photobleaching experiments in a droplet-by-droplet manner. To determine the residue length-dependent FRET efficiency within these condensates, the acceptor was photobleached, and a subsequent increase in donor intensity was recorded (Fig. 1l, m), which was used to extract ensemble FRET efficiency within individual condensates (Fig. 1n). The FRET efficiency for the N-to-108 construct was significantly higher than the N-to-86 and N-to-148 constructs. A higher FRET efficiency for the N-to-108 construct hinted at the presence of some long-range interactions in the polypeptide chain within these condensates. These ensemble FRET experiments are not capable of discerning conformational distribution and dynamics but provided the groundwork for carrying out more advanced single-molecule FRET measurements. Therefore, to detect and characterize the co-existing conformationally distinct subpopulations and their interconversion, we next set out to perform our single-molecule FRET measurements both in the dispersed and condensed phases.

### Experimental design for single-droplet single-molecule FRET

In this section, we provide a brief description of the experimental design for carrying out single-molecule experiments within individual condensates. A more detailed description of the setup, experiments, data acquisition, and data analysis are provided in Supplementary Information. Our single-molecule FRET experiments were performed using the two-color pulsed-interleaved excitation (PIE) mode on a time-resolved confocal microscope (Fig. 2). The PIE-FRET methodology permitted us to identify the low FRET states by eliminating the contribution of the zero-FRET efficiency peak which arises due to the acceptor photobleaching during the transit time^48^. In the case of the dispersed solutions, the dual-labeled protein concentration of 75-150 pM was sufficient to obtain single-molecule fluorescence bursts that arise due to freely diffusing fluorescently-labeled protein molecules within the femtolitre confocal volume. Photon bursts separated into the donor and acceptor channels provide the footprints of individual molecules that diffuse in and out of the confocal volume and allowed us to estimate the FRET efficiencies of individual diffusion events. In order to perform such single-molecule FRET experiments on individual condensates, a much lower concentration of the dual-labeled protein (5-10 pM) was needed in the condensed phase. Phase separation was induced from a mixture of unlabeled and dual-labeled FUS-LC so that the dual-labeled protein concentration was ∼ 5-10 pM and the total protein concentration was 200 μM. We placed the solution on a coverslip surface and allowed it to settle for ∼ 5 min after which most droplets got immobilized onto the glass surface. We then chose large immobilized droplets (3-7 µm diameter) that are much larger than the focal spot and focused the lasers inside these droplets (∼ 2 µm from the surface into the droplet). We used this procedure for single-droplet single-molecule FRET, FCS, and fluorescence anisotropy measurements. Fig. 2 shows representative time traces containing bursts in the dispersed phase and within single droplets. Using the PIE, only the bursts originating from the dual-labeled molecules qualify as FRET events for constructing FRET-efficiency histograms. The bursts originating from the droplets have longer duration compared to the monomeric protein which is attributed to the densely crowded environment and a slower diffusion within the condensed phase leading to a much larger number of excitation-emission cycles of the fluorophores during the transit time through the confocal volume. In the next two sections, we describe our single-molecule FRET results obtained in the monomeric and condensed phases of FUS-LC.

**Fig. 2.**
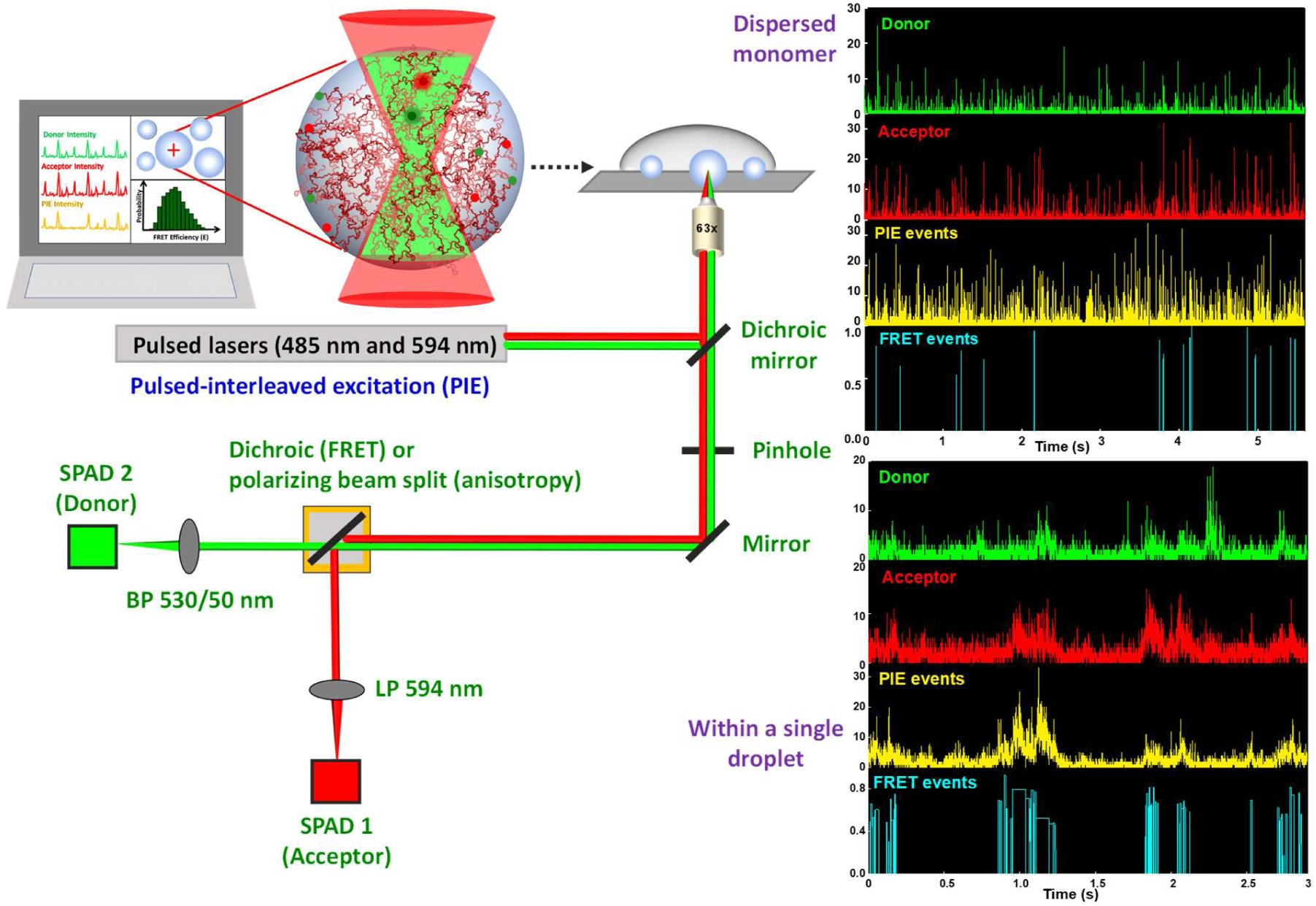
Experimental design for single-droplet single-molecule measurements. The schematic of our single-molecule microscopy setup (MicroTime 200, PicoQuant). The major components are an excitation system with two picosecond pulsed lasers (485 nm and 594 nm), an inverted microscope, and the confocal detection system consisting of an integrated system of dichroic mirrors, pinhole, bandpass filters, and single-photon avalanche diodes (SPADs) as detectors. Representative time traces displaying fluorescence bursts in donor and acceptor channels and corresponding PIE and FRET events recorded in the monomeric dispersed (top) and droplet (bottom) phases are also shown. See Supplementary Information for more details.

### Single-molecule FRET reveals two coexisting structural subpopulations of FUS-LC in the monomeric form

We performed our single-molecule FRET experiments with three dual-labeled constructs of FUS-LC varying the intramolecular distance (N-to-86, N-to-108, and N-to-148). Under the solution condition (pH 7.4, 250 mM NaCl), FUS-LC remains monomeric up to a concentration of 50 µM as evident by our dynamic light scattering measurements (Supplementary Fig. 1b, c). We performed single-molecule FRET experiments using 75-150 pM of dual-labeled FUS-LC. The single-molecule PIE-FRET efficiency histogram for the N-to-86 construct exhibited a unimodal distribution with a peak at ∼ 0.76 (Fig. 3a). The observed mean FRET efficiency was significantly higher than the calculated mean FRET efficiency expected for a coil in a good solvent (Supplementary Table 2) suggesting that the FUS-LC chain is considerably collapsed. Next, we chose two other constructs (N-to-108 and N-to-148) having larger intramolecular distances and expected much lower FRET efficiency. In contrast to our expectation, we observed an unusual bimodal distribution in the single-molecule FRET efficiency histogram for both constructs. The construct having a 108-residue separation (N-to-108) exhibited two FRET efficiency peaks at ∼ 0.98 and ∼ 0.80, whereas the N-to-148 construct showed peaks at ∼ 0.73 and ∼ 0.10 (Fig. 3b, c). These FRET efficiency histograms capture essential structural features indicating the presence of at least two predominant structural subpopulations in the monomeric conformational ensemble (Fig. 3d). These distinct subpopulations could involve compact paperclip-like (high-FRET) and partially extended tadpole-like (low-FRET) conformers that are in equilibrium having an interconversion exchange rate much slower than the observation time (0.5 ms). A binning time of 1 ms did not significantly alter the histograms suggesting the conformational exchange between these structural subpopulations could even be slower than 1 ms (Supplementary Fig. 2a, b). Such structural distributions can satisfactorily explain the observed chain length-dependent inter-residue FRET efficiency histograms. The major population comprises the high-FRET paperclip-like conformers that can potentially arise due to the strong intrachain interactions driven by π–π interactions between multiple tyrosine residues and hydrogen bonding between glutamine sidechains (Fig 1b). It is interesting to note that such compaction is observed in the N-terminal half of the polypeptide chain, whereas, the C-terminal segment adopts both compact and extended conformations possibly due to the presence of a larger number of proline residues. Taken together, our single-molecule FRET studies indicated that intrinsically disordered FUS-LC adopts two structurally distinct subpopulations having a varied extent of intrachain interactions. Next, we asked how conformational shapeshifting allows these intrachain interactions to turn into interchain interactions to promote phase separation of FUS-LC into liquid-like droplets.

**Fig. 3.**
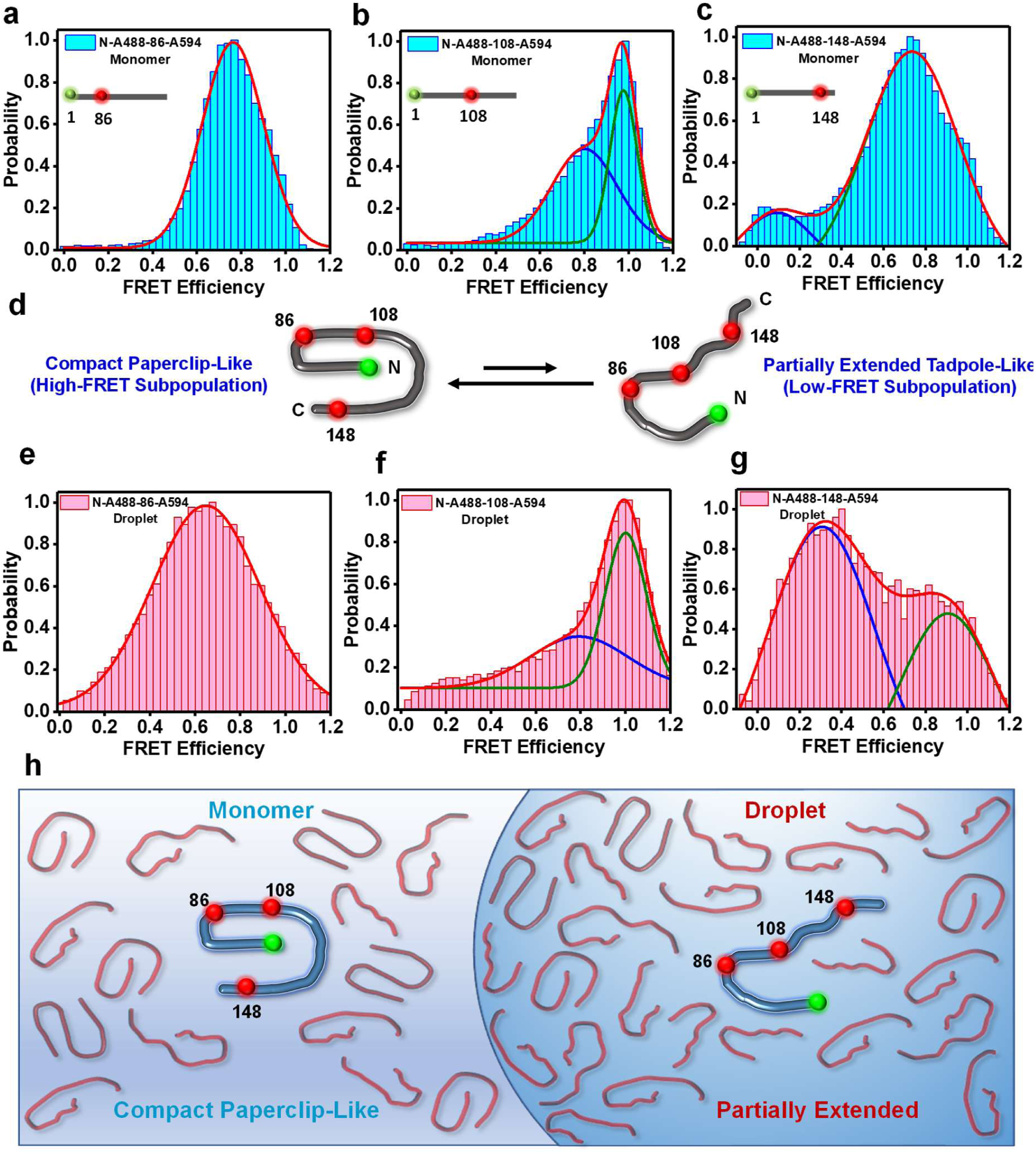
Single-molecule FRET in monomeric dispersed and droplet phases of FUS-LC. Single-molecule FRET histogram of monomeric FUS-LC in the monomeric dispersed phase for dual-labeled (a) N-to-86, (b) N-to-108, and (c) N-to-148 constructs. d. A schematic showing the coexistence of compact paperclip-like and extended tadpole-like conformers. (e-g) Single-droplet single-molecule FRET histograms for FUS-LC in condensed phase for dual-labeled N-to-86 (e), N-to-108 (f), and N-to-148 (g) constructs. The FRET efficiency values are shown in Supplementary Tables 2 and 3. The binning time was 0.5 ms and the total number of events was > 20,000. A binning time of 1 ms also yielded similar FRET histograms (Supplementary Fig. 2). See Supplementary Information for more details of experiments, data acquisition, and data analysis. h. A schematic depicting the structural unwinding of compact conformers into partially extended conformers upon phase separation.

### Single-droplet single-molecule FRET reveals a structural expansion and an increase in the conformational heterogeneity upon phase separation

We performed single-droplet single-molecule FRET measurements using all of the dual-labeled FUS-LC constructs (N-to-86, N-to-108, and N-to-148). The N-to-86 construct in the droplet phase also exhibited a unimodal FRET distribution with a lower mean FRET efficiency (∼0.64) (Fig. 3e) associated with a broader distribution compared to the dispersed monomeric form that showed a mean FRET efficiency of ∼ 0.76. This observation revealed the unwinding of the polypeptide chain that presumably allows the chains to participate in intermolecular interactions driving phase separation. The other two FRET constructs (N-to-108 and N-to-148) that showed bimodal FRET distribution in the monomeric form also exhibited a broadened distribution and a decrease in the energy transfer efficiencies upon phase separation (Fig. 3f, g). In the case of the N-to-108 construct, the mean FRET efficiencies of the two populations remained largely unaltered compared to the monomeric dispersed phase; however, the contribution of the low-FRET states grew with a concomitant broadening of the distribution. These results revealed that the N-terminal end and residues near the 108^th^ position are involved in long-range contacts that are persistent on the millisecond timescale. For the N-to-148 construct, the low-FRET states with a mean FRET efficiency of ∼ 0.30 constitute a major subpopulation within the droplet phase. It is interesting to note that these low-FRET states coexist with the high-FRET states in the condensed phase indicating conformational shapeshifting within the droplets occurs on a much slower timescale (>> 1 ms) than the typical observation time (Supplementary Fig. 2c, d). A shift in the FRET efficiency towards lower values accompanied by a growth in the lower efficiency population in the droplet phase signifies that partially extended conformers constitute the major subpopulation in the conformational ensemble along with the presence of compact paperclip-like conformers within the condensed phase. These findings revealed that the compact FUS-LC conformational ensemble undergoes considerable unwinding engendering more structural plasticity and heterogeneity that allow the dissolution of intramolecular interactions and the formation of new intermolecular contacts promoting phase separation (Fig. 3h). Glutamine and tyrosine residues in the condensed phase can participate in a dynamic network of intermolecular hydrogen bonding and π–π interactions within droplets giving rise to a highly condensed network fluid. These dynamic interactions can undergo making and breaking on a characteristic timescale giving rise to the internal viscoelastic behavior of these condensates. Therefore, next, we set out to study the polypeptide chain diffusion and dynamics within individual condensates on a wide range of timescales.

### Translational and rotational dynamics within individual condensates reveal the formation of a viscoelastic network fluid

In order to investigate the chain diffusion within individual condensates, we performed single-droplet FCS measurements using AlexaFluor488-labeled FUS-LC. FUS-LC exhibited a diffusion time of ∼ 0.13 ms in the monomeric dispersed form. The translational diffusion time increased 400 times to ∼ 50 ms in the droplet indicating a highly crowded and viscous environment within the droplets presumably due to the formation of a dynamic network of intermolecular interactions (Fig. 4a, b). We envisaged such a network of interactions giving rise to the condensate formation would impede the reorientation dynamics of polypeptide chains involved in intermolecular multivalent interactions. In order to delineate the role of reorientation dynamics, we employed site-specific single-droplet fluorescence (polarization) anisotropy measurements that report the extent of rotational flexibility of the polypeptide chain. For fluorescence anisotropy experiments, we chose four sites along the polypeptide chain and labeled single-Cys variants of FUS-LC (A16C, S86C, S108C, and S148C) using thiol-active fluorescein-5-maleimide that contains a shorter linker than Alexa dyes, and therefore, can report the rotational flexibility of the polypeptide chain without exhibiting a significant local depolarization. All residue locations exhibited a sharp increase in the steady-state fluorescence anisotropy in the condensed phase compared to the dispersed monomeric phase (Fig. 4c, d). These results indicated dampening of the rotational flexibility of the FUS-LC chain within droplets. Notably, the 108^th^ position exhibited a higher anisotropy value in both the monomer and droplet phases suggesting the possibility of some persistent long-range contacts that are in accordance with our single-molecule FRET data. Although steady-state fluorescence anisotropy measurements suggested reorientation restraints within condensates, these measurements do not allow us to discern the contributions of distinct modes of rotational dynamics. Next, we employed single-droplet picosecond time-resolved fluorescence anisotropy measurements that can discern the various modes of rotational dynamics.

**Fig. 4.**
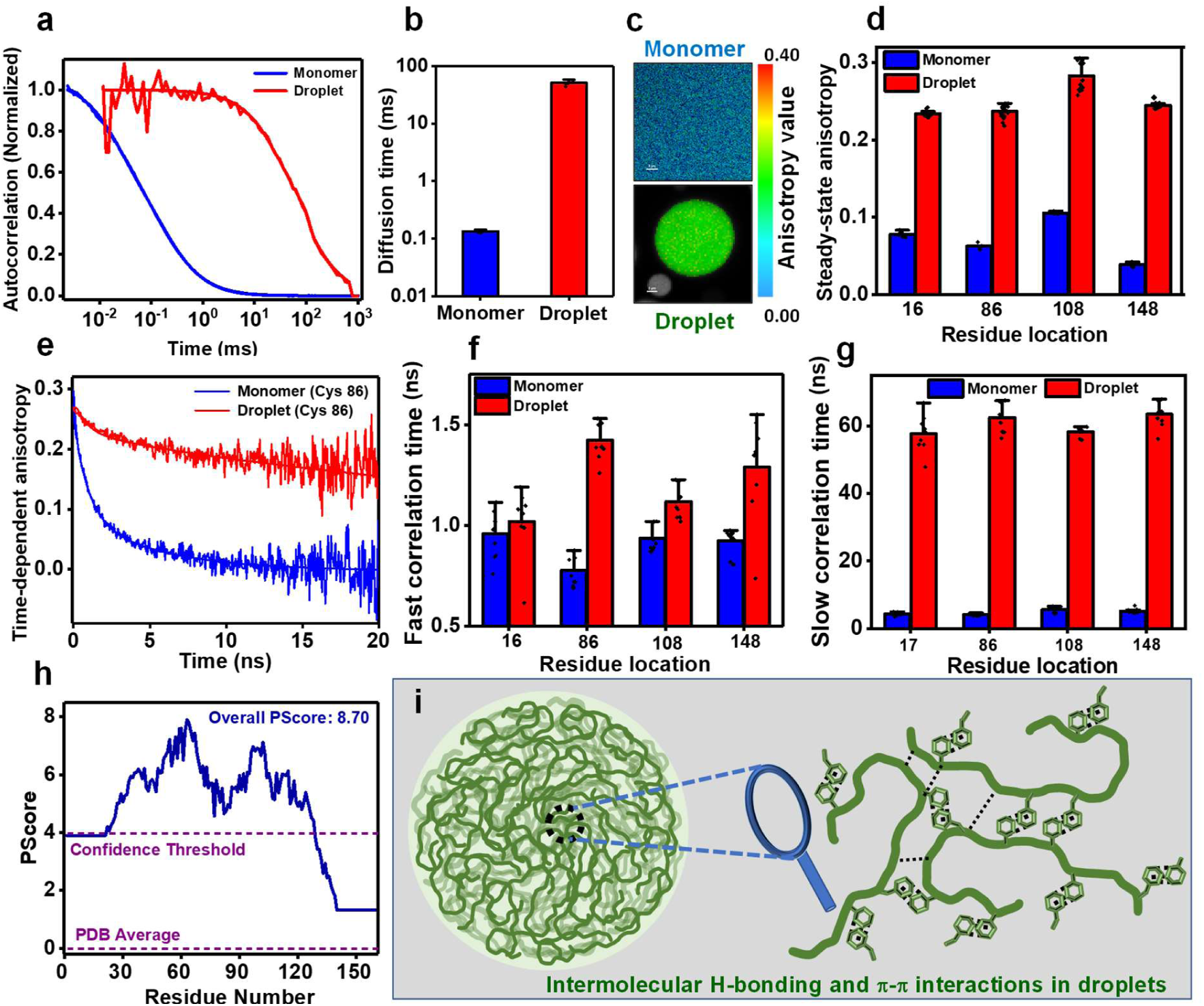
Translational and rotational dynamics. a. Autocorrelation plots (normalized) acquired from FCS measurements performed for AlexaFluor488 labeled FUS-LC in monomer and within single droplets (unnormalized FCS autocorrelation plots are shown in Supplementary Fig. 3a). b. The diffusion time of FUS-LC in monomeric and condensed phases estimated from FCS. Data represent mean ± SD for n = 5. FCS measurements were performed in the presence of 10 nM (in dispersed monomer) and 1-3 nM (in droplet) of AlexaFluor488-labeled FUS-LC. c. Representative fluorescence anisotropy images for a dispersed phase and a single droplet showing anisotropy heatmap. d. Steady-state fluorescence anisotropy values for FUS-LC in monomeric and within individual condensates. Fluorescence anisotropy was measured using fluorescein-5-maleimide-labeled FUS-LC at residue positions 16, 86, 108, and 148. Data represent mean ± SD for n ≥ 5 for monomers and n = 24 for droplets. e. Representative picosecond time-resolved fluorescence anisotropy decay profiles for FUS-LC labeled at residue position 86 in monomer and droplets. Solid lines are fits obtained from biexponential decay analysis. Time-resolved anisotropy decays for the other three locations (16, 108, and 148) are shown in Supplementary Fig. 3b-d. f. Fast rotational correlation times and g. slow rotational correlation times for residue locations 16, 86, 108, and 148 recovered from decay analyses. Data represent mean ± SD (n = 9). Rotational correlation times are included in Supplementary Table 4. h. Predictor of phase separation of IDPs based on the propensity to form long-range planar π-π contacts calculated as Pscore value for FUS-LC. i. A schematic representing a dense network of intermolecular π-π contacts and hydrogen bonding within FUS-LC condensates.

In order to temporally resolve the distinct molecular events, we utilized the highly sensitive picosecond time-resolved fluorescence anisotropy decay measurements that permit us to probe the depolarization kinetics of fluorescence anisotropy from the time-zero anisotropy value due to various modes of rotational relaxation^49^. In the case of monomeric IDPs, the depolarization kinetics follow a typical multiexponential decay function. Such anisotropy decay functions typically involve a fast rotational correlation time representing the local wobbling-in-cone motion of the fluorophore and slow rotational correlation times corresponding to backbone dihedral angle fluctuations and long-range reorientation dynamics^50–52^. As expected, the anisotropy decay profiles for monomeric FUS-LC recorded at different residue locations were satisfactorily described by a biexponential decay model giving rise to two well-separated correlation times: fast rotational correlation time representing a local probe motion (∼ 1 ns) and a slow rotational correlation time (∼ 4-5 ns) corresponding to dihedral and long-range reorientation motions (Fig. 4e-g). We then measured fluorescence anisotropy decay kinetics within individual droplets and observed that the depolarization kinetics slowed down considerably. The fast local correlation time exhibited a slight increase, whereas, the slow rotational correlation displayed a sharp increase from ∼ 5 ns to ∼ 60 ns at all residue locations (Fig. 4e-g). Such an increase in the slow correlation time within condensates indicated a dampening of the chain reorientation dynamics presumably due to the formation of a network via interchain physical crosslinks. From the amino acid composition, we postulated that interchain contacts are possibly formed primarily via aromatic π–π and hydrogen bonding interactions between tyrosine and glutamine residues, respectively^24^ (Fig. 1b). The PScore analysis^18^ that quantifies the π–π contacts in proteins revealed a high propensity of π–π interactions mediated phase separation of FUS-LC which contains 24 tyrosine and 37 glutamine residues (Fig. 4h) (Supplementary Fig. 3e). Taken together, our results on chain dynamics coupled with structural subpopulations indicate the presence of a network of intermolecular interactions as depicted in our schematic (Fig. 4i). Such a network of dynamic physical crosslinks can slow down translational and rotational diffusion ensuing a viscoelastic network fluid within the condensates. Next, we asked if our unique structural and dynamical readouts of wild-type FUS-LC can detect and distinguish the altered phase behavior of a disease-associated mutation.

### A disease-associated mutant alters the phase behavior by modifying conformational distribution and dynamics

Several mutations in the disordered LC domain, RNA-binding domain, and NLS are associated with various neurodegenerative diseases. One such mutation in the LC domain (G156E) is a patient-derived, clinically relevant mutation that functions by modulating the phase behavior and aggregation propensity of FUS^33, 53, 54^. Thus, we next set out to study the effect of this disease-related mutation on the conformational characteristics and phase behavior of FUS-LC. We created a single-point mutant (G156E) and recombinantly expressed this construct. The CD spectrum of G156E FUS-LC indicated a disordered conformation and showed no significant changes in the secondary structural contents as compared to the wild-type FUS-LC (Supplementary Fig. 4a). Next, we began with the phase separation assay of G156E FUS-LC. A rise in turbidity values of the protein solution in the presence of salt (Supplementary Fig. 4b) indicated the formation of droplets capable of recruiting dual-labeled wild-type FUS-LC as confirmed by our two-color confocal fluorescence imaging (Fig. 5a). As predicted by the phase separation propensity prediction tool catGranule^55^ (Supplementary Fig. 4c), the phase separation propensity of G156E FUS-LC was slightly lower in comparison to the wild-type FUS-LC as indicated by a relatively lower turbidity (Supplementary Fig. 1a, 4b) and a higher saturation concentration compared to wild-type FUS-LC (Fig. 5b). To further characterize these G156E FUS-LC droplets, we performed FRAP measurements which showed a slower and lower fluorescence recovery compared to wild-type FUS-LC (Fig. 5c). In order to get insights into the polypeptide chain diffusion within individual droplets, we performed single-droplet FCS measurements which indicated a slightly slower translational diffusion (∼ 70 ms) in comparison to wild-type FUS-LC droplets (∼ 50 ms) (Fig. 5d, Supplementary Fig. 4d). These results suggested that the G156E mutation alters the phase behavior and the condensed phase exhibits more restrained diffusion.

**Fig. 5.**
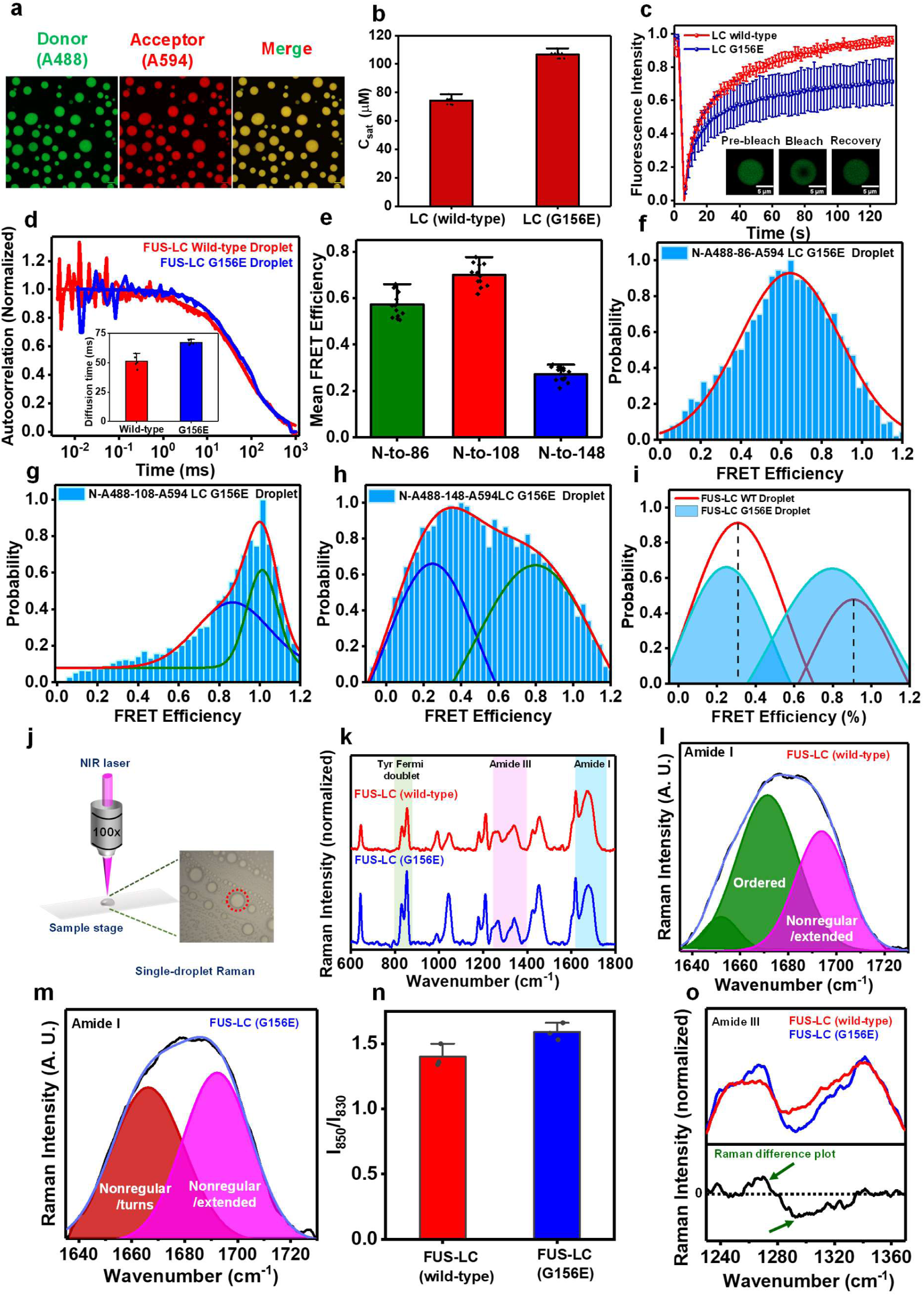
Effect of disease-associate mutation (G156E) on FUS-LC. a. Airy scan confocal imaging showing G156E FUS-LC droplets recruit wild-type dual-labeled FUS-LC. b. Saturation concentrations (C_sat_) of wild-type and G156E FUS-LC were estimated using centrifugation. Data represent mean ± SD (n = 9). c. FRAP kinetics of AlexaFluor488-labeled FUS-LC within wild-type and G156E mutant droplets. Data represent mean ± SD (n = 8). The FRAP kinetics for wild-type FUS-LC (red) is the same as in Fig. 1i and are included here for comparison. d. Normalized FCS autocorrelation plots for wild-type and G156E droplets. Inset shows the mean diffusion times for droplets of wild-type and G156E FUS-LC. Data represent mean ± SD (n = 5). Data for wild-type FUS-LC droplets are the same in Fig. 4a,b and included here for comparison. e. Mean FRET efficiency obtained from acceptor photobleaching of G156E FUS-LC droplets containing 0.05% of dual-labeled FUS-LC. Data represent mean ± SD (n = 13). Single-droplet single-molecule FRET histograms in G156E FUS-LC droplets using dual-labeled (f) N-to-86, (g) N-to-108, and (h) N-to-148 constructs. i. Overlay of fitted subpopulations for wild-type and G156E FUS-LC droplets (shaded) for the N-to-148 construct showing a shift towards a lower FRET efficiency within G156E FUS-LC droplets. The binning time was 0.5 ms and the total number of events was > 20,000. FRET efficiency values are shown in Supplementary Table 3. j. A schematic for our single-droplet Raman by focusing a near-infrared (NIR) laser using a 100x objective into a single droplet of FUS-LC. k. Raman spectra of individual droplets of wild-type and G156E FUS-LC (n = 3). Raman spectra of three individual droplets are shown in Supplementary Fig. 5. Raman spectra are normalized with respect to the amide I band at ∼ 1673 cm^-1^. l,m. Gaussian deconvolution of amide I band of wild-type FUS-LC (l) and G156E FUS-LC (m) to estimate the percentage composition of various secondary structural elements. The black and blue solid lines represent the actual data and the cumulative fit, respectively. The colored solid lines represent the Gaussian peaks obtained after deconvolution. n. The ratio of the tyrosine Fermi doublet (I_850_/I_830_) estimated from the peak intensities. The mean ratio and standard error are estimated from at least three independent measurements. o. The Raman difference plot of the amide III band of FUS-LC droplets and G156E FUS-LC droplets (arrow indicates the difference of interest). See Supplementary Information for more experimental details.

Next, in order to gain structural insights into the wild-type FUS-LC chain within G156E FUS-LC condensates, we performed ensemble single-droplet FRET using acceptor photobleaching in the confocal fluorescence microscopy format. Ensemble single-droplet FRET indicated the retention of long-range contacts between the N-terminal and residue location 108 similar to the wild-type FUS-LC droplets (Fig. 5e). To obtain insights into the structural subpopulations and conformational dynamics, we then performed single-droplet single-molecule FRET measurements (Fig. 5f-h). FRET efficiency histograms for the N-to-86 and N-to-108 constructs within G156E droplets were similar to those observed for wild-type FUS-LC droplets. Interestingly, the FRET efficiency histogram for the N-to-148 construct exhibited a slight shift towards lower FRET efficiency compared to the wild-type FUS-LC droplets (Fig. 5i). This reduction in FRET efficiency of both populations indicated more unwinding of the C-terminal segment of the chain that facilitates the formation of a network of intermolecular contacts within the G156E FUS-LC condensates. Such a conformational unwinding in the mutant, presumably due to the presence of an additional negative charge, can offer a larger number of multivalent interactions leading to a more densely packed interior of the condensate and more dampened translational and rotational dynamics. This observation is consistent with a slight increase in the steady-state fluorescence anisotropy within G156E FUS-LC condensates (Supplementary Fig. 4e) and a slower internal diffusion corroborating our FRAP and FCS results. Taken together, our results revealed an unraveling of the polypeptide chain with condensates of the G156E mutant promoting increased interchain interactions and a dense network which can further facilitate pathological aggregation of the mutant FUS-LC as observed in previous studies^33, 53, 54^. Next, we wanted to distinguish the secondary structural features that govern the condensate properties of wild-type and mutant FUS-LC.

### Single-droplet vibrational Raman spectroscopy supports the altered phase behavior of the disease-associated mutant

In order to probe the secondary structural features of FUS-LC, we performed single-droplet vibrational Raman spectroscopy that allows us to obtain insights into polypeptide structure and organization within individual droplets^56, 57^. Focusing a laser beam into a droplet permits us to obtain a Raman spectrum that contains the characteristic vibrational signatures corresponding to the polypeptide backbone (amide I and amide III) and side chains (aromatic and aliphatic residues). These vibrational signatures provide us with vital information on the secondary structural distributions and intermolecular interactions within a single condensate (Fig. 5j). Amide I vibrational band (1630-1700 cm^-1^) originates primarily due to the C=O stretching vibrations of the polypeptide backbone, while the amide III band (1230-1320 cm^-1^) involves a combination of C-N stretch and N-H bending motions of the backbone. Together these amide bands constitute the secondary structural marker bands highlighting the secondary structural elements present in the proteins^24, 56–59^. We observed a broad amide I band for wild-type FUS-LC droplets indicating the presence of disordered conformations and considerable conformational heterogeneity in the condensed phase (Fig. 5k) corroborating our single-molecule FRET results. Deconvolution of amide I indicated the presence of two major peaks representing extended/unordered (∼ 1692 cm^-1^) and some ordered structural elements in the protein-rich dense phase. The peak at ∼ 1671 cm^-1^ corresponds to ordered structural elements that might contain β-sheet with some minor α-helical contents (∼ 1652 cm^-1^) and can represent the signature of compact paperclip-like states as observed in our FRET experiments (Fig. 5l). On the contrary, the deconvolution analysis of the amide peak for G156E FUS-LC droplets revealed a largely unstructured state (∼ 1666 cm^-1^ and ∼ 1692 cm^-1^) within droplets corroborating our FRET data (Fig. 5m). A small increase in the full width at half maximum (FWHM) of the amide I band from wild-type (49.6 ± 0.6 cm^-1^) to mutant droplets (51.4 ± 0.6 cm^-1^) could further support more conformational heterogeneity in the case of the G156E mutant. More structural unzipping and heterogeneity in the case of the mutant allows a much denser network of interaction as observed in our single-molecule FRET, FCS, and anisotropy measurements. This is also supported by the Raman intensity ratio of the tyrosine Fermi doublet (I_850_/I_830_), which indicates the hydrogen bonding propensity of the phenolic hydroxyl group of tyrosine with surrounding water molecules or in other words, determines the average solvent accessibility of tyrosine residues. This ratio changed from ∼ 1.4 for wild-type droplets to ∼ 1.6 for mutant droplets indicating a higher solvent exposure of tyrosine residues within G156E FUS-LC droplets resulting from relatively expanded conformers facilitating a larger extent of intermolecular π-π contacts between more expanded chains within G156E FUS-LC droplets (Fig. 5n). We further zoomed into the amide III region for wild-type and G156E droplets and constructed a Raman difference plot for the comparison. A positive band centered at ∼ 1270 cm^-1^ (nonregular/turns) and a negative band at ∼ 1300 cm^-1^ highlighted a higher content of disordered conformations within mutant droplets as compared to wild-type droplets (Fig. 5o). Our single-droplet vibrational Raman studies capture the key structural differences between wild-type and mutant FUS-LC droplets and are in agreement our single-molecule FRET, FCS, FRAP, and anisotropy results. Taken together, the disease-associated mutant (G156E) of FUS-LC exhibits higher disorder facilitating more intermolecular association with altered material properties that can potentially promote aberrant phase transitions into solid-like pathological aggregates compared to the wild-type form.

## Discussion

In this work, we showed that the polypeptide chain of FUS-LC undergoes conformational shapeshifting from the monomeric dispersed phase to the condensed phase. We summarized all our observations in a schematic illustration in Figure 6. By employing single-molecule FRET, we characterized the conformational distribution and dynamics of FUS-LC, site-specifically and orthogonally labeled with donor and acceptor fluorophores. Such single-molecule studies carried out at ultralow concentrations (∼ 100 pM) allow us to unambiguously interrogate and characterize the monomeric form of the protein in a molecule-by-molecule manner that is not generally achievable by most other conventional structural tools. Our single-molecule FRET studies revealed that intrinsically disordered FUS-LC monomer exists as a heterogeneous ensemble of structures comprising chiefly two distinct, well-resolved, structural subpopulations. These subpopulations include compact paperclip-like conformers and partially extended tadpole-like conformers exchanging on a much slower timescale (>> 1 ms) than the observation time. The compaction of the disordered FUS-LC chain can be ascribed to the presence of extensive intramolecular interactions arising due to π–π interactions between multiple tyrosine residues and hydrogen bonding between glutamine side chains. We propose that a large number of proline and glycine residues promote partial expansion, especially at the C-terminal part of the polypeptide chain. Such making and breaking of intramolecular noncovalent contacts give rise to rapidly fluctuating, intrinsically disordered, heterogeneous conformational ensemble in the monomeric form of FUS-LC. This conformational equilibrium in the infinitesimally dilute condition changes quite dramatically when the protein concentration is raised beyond a threshold concentration also known as the saturation concentration. Above this concentration, intermolecular chain-chain interactions become more dominant than intramolecular contacts, and thus, unzipping of compact conformers becomes facile due to the more favorable intermolecular multivalent interactions driving liquid phase condensation. The slow conformational exchange between the conformers allows more interaction lifetime yielding a network of intermolecular interactions in the condensed phase. Therefore, intriguing conformational gymnastics play a pivotal role in driving protein phase transitions.

**Fig. 6.**
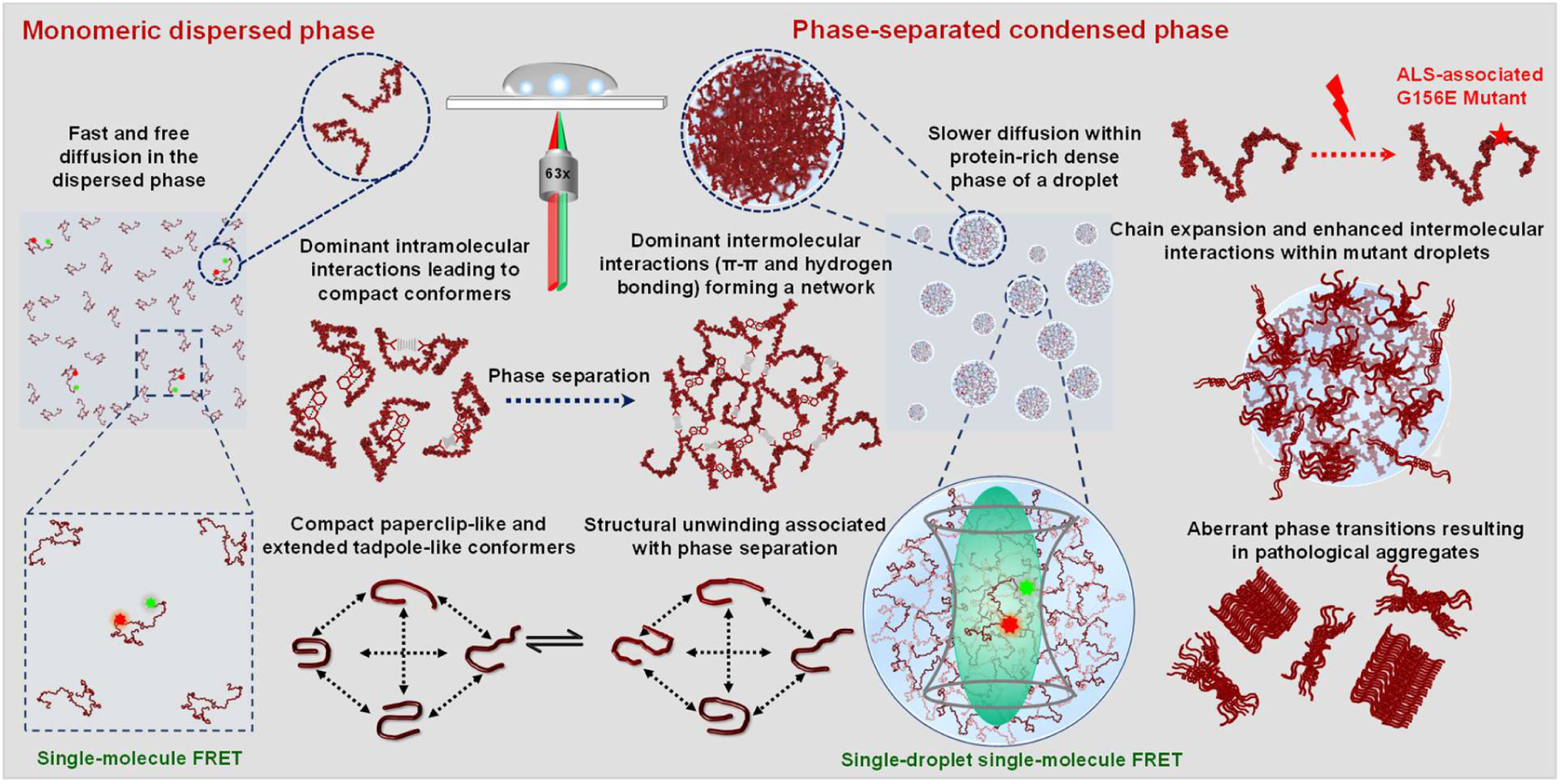
A schematic of conformational shapeshifting during phase separation. A summary of our single-molecule FRET, FCS, fluorescence anisotropy, Raman spectroscopy experiments, and the impact of a pathological mutation in the phase behavior of FUS-LC.

Indeed, our single-droplet single-molecule FRET studies revealed a symphony of conformational shapeshifting events within the dense phase of individual condensates as depicted in Figure 6. Some of these unique molecular features involving distinct subpopulations generally remain skewed in traditional bulk experiments and can only be directly deciphered using single-molecule tools. Single-molecule FRET coupled with FCS and picosecond time-resolved fluorescence anisotropy measurements allowed us to directly probe the conformational distribution and dynamics within the dense phase. The solvent quality within this protein-rich dense phase is likely to be better than that in the monomeric form of the protein dispersed in water which is a poor solvent for a polypeptide chain. A better solvent quality, and hence a higher Flory scaling exponent, in the dense phase is expected to favor the conformational expansion. Such expanded conformers can also turn intramolecular contacts into transient intermolecular contacts favoring the condensed phase. The presence of a subpopulation of compact states suggests that the conformational interconversion is possibly associated with the making and breaking of multivalent interactions giving rise to liquid-like behavior within droplets. An interesting hypothesis can be posited based on the radial gradient of the physicochemical properties with a complex interplay of intra-chain, inter-chain, and chain-solvent interactions in the condensed phase^60^. These droplets could possess a radial distribution of conformational subpopulations based on the spatial locations within the condensate. Previous simulation studies on intrinsically disordered prion-like low-complexity domains have indicated such conformational heterogeneity within the dense phase ^60, 61^. Recent studies revealed that a small-world percolated network comprising a varied density of physical crosslinks can give rise to distinct conformational states and molecular orientations varying spatial location from the center to the interface of the condensates^60^. The spatial resolution of the confocal microscopy-based format used in our single-molecule experiments is inadequate for dissecting the radial distribution of conformers. The FRET efficiency distribution displayed by our observed single-molecule FRET histograms could potentially comprise such conformational distributions across the droplet locations. We would also like to point out that although we observed much broader peaks in the FRET efficiency histograms in droplets, we are unable to comment on the width of individual peaks since there can be some contributions from the shot noise due to low photon counts and the orientation factor that can potentially contribute to the widths of the distribution. Our single-droplet FCS and fluorescence anisotropy results revealed that both translational diffusion and rotational chain reorientation dynamics are considerably slowed down within condensates. These findings hinted at the formation of a network of physical crosslinks giving rise to viscoelastic network fluid. Our results on a disease-associated mutant (G156E) indicated a more conformational plasticity promoting a denser network of intermolecular contacts as observed by single-molecule FRET, FRAP, FCS, and anisotropy measurements. Additionally, single-droplet vibrational Raman spectroscopy corroborated our results on altered conformational distribution for the mutant. Raman signatures for both backbone and sidechain markers indicated slightly more extended conformers and greater participation in chain-chain association within condensates. Such interactions can potentially promote aberrant liquid-to-solid phase transitions and accelerated aggregation associated with the pathological hallmark of the G156E mutant of FUS (Fig. 6).

In summary, our single-molecule experiments directly unveiled an intriguing interplay of conformational heterogeneity, structural distribution, and dynamics that crucially governs the course of phase transitions of prion-like low-complexity domains. Such low-complexity domains are present in FUS and other FUS-family proteins and are essential in mediating the homotypic and heterotypic interactions driving the assembly into liquid-like functional condensates and solid-like pathological aggregates. Post-translational modifications and mutations can alter the interplay between intra- and intermolecular interactions shifting the conformational equilibria both in the dispersed and condensed phases. Such altered interactions can give rise to changes in the viscoelastic material properties of biomolecular condensates promoting aberrant phase transitions associated with ALS and FTD. Our findings on the role of conformational excursions in phase separation can have much broader implications for a wide range of phase-separating proteins involved in physiology and disease. For instance, tau, an intrinsically disordered neuronal protein associated with Alzheimer’s disease, exhibits conformational subpopulations comprising paperclip and expanded states^41, 62–64^. Phase separation of tau studied at the single-molecule resolution also revealed an increased conformational heterogeneity. We suggest that the sequence-encoded structural unwinding coupled with a dynamic control can expose the multivalency of polypeptide chains that promote ephemeral interactions resulting in biomolecular condensate formation. Deriving general principles from single-molecule studies on a wide range of phase-separating proteins and artificial polypeptides can provide the key mechanistic underpinning of macromolecular phase separation and pave the way for novel synthetic biology applications. Additionally, the development of multi-color, multi-parameter, super-resolved, single-droplet single-molecule FRET can offer unprecedented spatiotemporal resolution in studying intracellular heterotypic, multicomponent, and multiphasic biomolecular condensates and for exploring unchartered territories of biological phase transitions.

## Methods

FUS-LC variants were recombinantly expressed and purified using affinity and size exclusion chromatography. Airy scan confocal imaging and FRAP experiments were performed on a ZEISS LSM 980 Elyra 7 super-resolution microscope using a 63x oil-immersion objective (N.A. = 1.4). Single-molecule FRET and FCS experiments were performed using a MicroTime 200 time-resolved confocal microscope (PicoQuant). For single-molecule FRET, single-cysteine mutants were dual-labeled with amine and thiol-reactive FRET dyes: AlexaFluor488 NHS (for N-terminal labeling) and AlexaFluor594-maleimide (for Cys labeling). The conformational distributions and heterogeneity were studied using single-molecule FRET experiments within monomeric dispersed and droplet phases. The translational diffusion was measured using single-droplet FCS. Steady-state fluorescence anisotropy imaging and time-resolved fluorescence anisotropy decay measurements were made on the MicroTime 200 time-resolved confocal microscope. Single-droplet vibrational Raman scattering was recorded using an inVia laser Raman microscope (Renishaw, UK). See Supplementary Information for experimental and data analyses.

## Acknowledgments

We thank IISER Mohali, Science and Engineering Research Board, (SUPRA SPR/2020/000333 to S.M.) and Department of Science and Technology Ministry of Education, Govt. of India for (FIST grant # SR/FST/LS-II/2017/97 to the Department of Biological Sciences, IISER Mohali and Centre of Excellence grant to S.M), for financial support. Prof. N. Periasamy (Retd. TIFR Mumbai) for the fluorescence decay analysis program. We thank Swastik G. Pattanashetty for his valuable contribution during the initial phase of the project. Dr. Mily Bhattacharya and current and former members of Mukhopadhyay Lab for their valuable input and for critically reading this manuscript.

## Authors contributions

A.J. and S.M. conceived the project. A.J., A.W., and S.M. further developed the concept and the experimental design. A.J., A.W., A.A., S.K.R., L.A., and S.S. performed the experiments and analyses. A.J. prepared the figures and wrote the first draft. S.M. supervised the work, wrote/edited the manuscript, obtained funding, and provided the overall direction. All authors discussed the results and commented on the manuscript.

## References

1. Kilgore, H. R. & Young, R. A. Learning the chemical grammar of biomolecular condensates. Nat. Chem. Biol. 18, 1298–1306 (2022).

2. Alberti, S. & Hyman, A. A. Biomolecular condensates at the nexus of cellular stress, protein aggregation disease and ageing. Nat. Rev. Mol. Cell Biol. 22, 196–213 (2021).

3. Lyon, A. S., Peeples, W. B., & Rosen, M. K. A framework for understanding the functions of biomolecular condensates across scales. Nat. Rev. Mol. Cell Biol. 22, 215–235 (2021).

4. Mittag, T. & Pappu, R. V. A conceptual framework for understanding phase separation and addressing open questions and challenges. Mol. Cell 82, 2201–2214 (2022).

5. Boeynaems, S., Alberti, S., Fawzi, N. L., Mittag, T., Polymenidou, M., Rousseau, F., Schymkowitz, J., Shorter, J., Wolozin, B., Van Den Bosch, L., Tompa, P., & Fuxreiter, M. Protein Phase Separation: A New Phase in Cell Biology. Trends Cell Biol. 28, 420–435 (2018).

6. Roden, C., & Gladfelter, A. S. RNA contributions to the form and function of biomolecular condensates. Nat. Rev. Mol. Cell Biol. 22, 183–195 (2021).

7. Choi, J. M., Holehouse, A. S., & Pappu, R. V. Physical principles underlying the complex biology of intracellular phase transitions. Annu. Rev. Biophys. 49, 107–133 (2020).

8. Dignon, G. L., Best, R. B., & Mittal, J. Biomolecular phase separation: From molecular driving forces to macroscopic properties. Annu. Rev. Phys. Chem. 71, 53–75 (2020).

9. Yu, M., Heidari, M., Mikhaleva, S., Tan, P. S., Mingu, S., Ruan, H., Reinkemeier, C. D., Obarska-Kosinska, A, Siggel, M., Beck, M., Hummer, G., & Lemke, E. A. Visualizing the disordered nuclear transport machinery in situ. Nature 617, 162–169 (2023).

10. Shapiro, D. M., Ney, M., Eghtesadi, S. A., & Chilkoti, A. Protein phase separation arising from intrinsic disorder: First-principles to bespoke applications. J. Phys. Chem. B 125, 6740– 6759 (2021).

11. Mitrea, D. M., Cika, J. A., Guy, C. S., Ban, D., Banerjee, P. R., Stanley, C. B., Nourse, A., Deniz, A. A., & Kriwacki, R. W. Nucleophosmin integrates within the nucleolus via multi-modal interactions with proteins displaying R-rich linear motifs and rRNA. eLife 5, (2016).

12. Shin, Y. & Brangwynne, C. P. Liquid phase condensation in cell physiology and disease. Science 357, (2017).

13. Mitrea, D. M., & Kriwacki, R. W. Phase separation in biology; functional organization of a higher order. Cell Commun. Signal. 14, (2016).

14. Martin, E. W., Holehouse, A. S., Peran, I., Farag, M., Incicco, J. J., Bremer, A., Grace, C. R., Soranno, A., Pappu, R. V., & Mittag, T. Valence and patterning of aromatic residues determine the phase behavior of prion-like domains. Science 367, 694–699 (2020).

15. Mukhopadhyay, S. The dynamism of intrinsically disordered proteins: Binding induced folding, amyloid formation, and phase separation. J. Phys. Chem. B 124, 11541–11560 (2020).

16. Martin, E. W. & Mittag, T. Relationship of sequence and phase separation in protein low-complexity regions. Biochemistry 57, 2478–2487 (2018).

17. Brangwynne, C. P., Tompa, P., & Pappu, R. V. Polymer physics of intracellular phase transitions. Nat. Phys. 11, 899–904 (2015).

18. Vernon, R. M., Chong, P. A., Tsang, B., Kim, T. H., Bah, A., Farber, P., Lin, H., & Forman-Kay, J. D. Pi-Pi contacts are an overlooked protein feature relevant to phase separation. eLife 9, (2018).

19. Ruff, K. M., Choi, Y. H., Cox, D., Ormsby, A. R., Myung, Y., Ascher, D. B., Radford, S. E., Pappu, R. V., & Hatters, D. M. Sequence grammar underlying the unfolding and phase separation of globular proteins. Mol. Cell. 82, 3193–3208 (2022).

20. Uversky, V. N. Intrinsically disordered proteins in overcrowded milieu: Membrane-less organelles, phase separation, and intrinsic disorder. Curr. Opin. Struct. Biol. 44, 18–30 (2017).

21. Kaur, T., Raju, M., Alshareedah, I., Davis, R.B., Potoyan, D.A., & Banerjee, P. R. Sequence-encoded and composition-dependent protein-RNA interactions control multiphasic condensate morphologies. Nat. Commun. 12, (2021).

22. Vendruscolo, M., & Fuxreiter, M. Protein condensation diseases: therapeutic opportunities. Nat. Commun. 13, (2022).

23. Svetoni, F., Frisone, P. & Paronetto, M. P. Role of FET proteins in neurodegenerative disorders. RNA Biol. 13, 1089–1102 (2016).

24. Murthy, A. C., Dignon, G. L., Kan, Y., Zerze, G. H., Parekh, S. H., Mittal, J., & Fawzi, N. L. (2019). Molecular interactions underlying liquid-liquid phase separation of the FUS low-complexity domain. Nat. Struct. Mol. Biol. 26, 637–648 (2019).

25. Loughlin, F. E., Lukavsky, P. J., Kazeeva, T., Reber, S., Hock, E. M., Colombo, M., Von Schroetter, C., Pauli, P., Cléry, A., Mühlemann, O., Polymenidou, M., Ruepp, M. D., & Allain, F. H. The Solution Structure of FUS Bound to RNA Reveals a Bipartite Mode of RNA Recognition with Both Sequence and Shape Specificity. Mol. Cell 73, 490–504 (2019).

26. Hofweber, M., Hutten, S., Bourgeois, B., Spreitzer, E., Niedner-Boblenz, A., Schifferer, M., Ruepp, M. D., Simons, M., Niessing, D., Madl, T., & Dormann, D. Phase Separation of FUS Is Suppressed by Its Nuclear Import Receptor and Arginine Methylation. Cell 173, 706–719 (2018).

27. Qamar, S., Wang, G., Randle, S. J., Ruggeri, F. S., Varela, J. A., Lin, J. Q., Phillips, E. C., Miyashita, A., Williams, D., Ströhl, F., Meadows, W., Ferry, R., Dardov, V. J., Tartaglia, G. G., Farrer, L. A., Kaminski Schierle, G. S., Kaminski, C. F., Holt, C. E., Fraser, P. E., Schmitt-Ulms, G., Klenerman, D., Knowles, T., Vendruscolo, M., & St George-Hyslop, P. FUS Phase Separation Is Modulated by a Molecular Chaperone and Methylation of Arginine Cation-π Interactions. Cell 173, 720–734 (2018).

28. Murray, D. T., Kato, M., Lin, Y., Thurber, K.R., Hung, I., McKnight, S. L., & Tycko, R. Structure of FUS Protein Fibrils and Its Relevance to Self-Assembly and Phase Separation of Low-Complexity Domains. Cell. 171, 615–627 (2017).

29. Burke, K. A., Janke, A. M., Rhine, C. L., & Fawzi, N. L. Residue-by-Residue View of In Vitro FUS Granules that Bind the C-Terminal Domain of RNA Polymerase II. Mol. Cell 15, 231–41(2015).

30. Murthy, A. C., Tang, W. S., Jovic, N., Janke, A. M., Seo, D. H., Perdikari, T. M., Mittal, J., & Fawzi, N. L. Molecular interactions contributing to FUS SYGQ LC-RGG phase separation and co-partitioning with RNA polymerase II heptads. Nat. Struct. Mol. Biol. 28, 923–935 (2021).

31. Monahan, Z., Ryan, V. H., Janke, A. M., Burke, K. A., Rhoads, S. N., Zerze, G. H., O’Meally, R., Dignon, G. L., Conicella, A. E., Zheng, W., Best, R. B., Cole, R. N., Mittal, J., Shewmaker, F., & Fawzi, N. L. Phosphorylation of the FUS low-complexity domain disrupts phase separation, aggregation, and toxicity. EMBO J. 36, 2951–2967 (2017).

32. Lee, M., Ghosh, U., Thurber, K. R., Kato, M., & Tycko, R. Molecular structure and interactions within amyloid-like fibrils formed by a low-complexity protein sequence from FUS. Nat. Commun. 11, (2020).

33. Berkeley, R. F., Kashefi, M., & Debelouchina, G. T. Real-time observation of structure and dynamics during the liquid-to-solid transition of FUS LC. Biophys. J. 120, 1276–1287 (2021).

34. Emmanouilidis, L., Esteban-Hofer, L., Damberger, F. F., de Vries, T., Nguyen, C. K. X., Ibáñez, L. F., Mergenthal, S., Klotzsch, E., Yulikov, M., Jeschke, G., & Allain, F. H. NMR and EPR reveal a compaction of the RNA-binding protein FUS upon droplet formation. Nat. Chem. Biol. 17, 608–614 (2021).

35. Kato, M., & McKnight, S. L. The low-complexity domain of the FUS RNA binding protein self-assembles via the mutually exclusive use of two distinct cross-β cores. Proc. Natl. Acad. Sci. USA 19, (2021).

36. Schuler, B., & Hofmann, H. Single-molecule spectroscopy of protein folding dynamics--expanding scope and timescales. Curr. Opin. Struct. Biol. 23, 36–47 (2013).

37. Agam, G., Gebhardt, C., Popara, M., Mächtel, R., Folz, J., Ambrose, B., Chamachi, N., Chung, S. Y., Craggs, T. D., de Boer, M., Grohmann, D., Ha, T., Hartmann, A., Hendrix, J., Hirschfeld, V., Hübner, C. G., Hugel, T., Kammerer, D., Kang, H. S., Kapanidis, A. N., Krainer, G., Kramm, K., Lemke, E. A., Lerner, E., Margeat, E., Martens, K., Michaelis, J., Mitra, J., Moya Muñoz, G. G., Quast, R. B., Robb, N. C., Sattler, M., Schlierf, M., Schneider, J., Schröder, T., Sefer, A., Tan, P. S., Thurn, J., Tinnefeld, P., van Noort, J., Weiss, S., Wendler, N., Zijlstra, N., Barth, A., Seidel, C. A. M., Lamb, D. C., & Cordes, T. Reliability and accuracy of single-molecule FRET studies for characterization of structural dynamics and distances in proteins. Nat. Methods. 20, 523–535 (2023).

38. Brucale, M., Schuler, B., & Samorì, B. Single-molecule studies of intrinsically disordered proteins. Chem. Rev. 26, 3281–317 (2014).

39. Nasir, I., Bentley, E. P., & Deniz, A. A. Ratiometric Single-Molecule FRET Measurements to Probe Conformational Subpopulations of Intrinsically Disordered Proteins. Curr. Protoc. Chem. Biol. 12, (2020).

40. Feng, X. A., Poyton, M. F., & Ha, T. Multicolor single-molecule FRET for DNA and RNA processes. Curr. Opin. Struct. Biol. 70, 26–33 (2021).

41. Elbaum-Garfinkle, S., & Rhoades, E. Identification of an aggregation-prone structure of tau. J. Am. Chem. Soc. 10, 16607–13 (2012).

42. Galvanetto, N., Ivanović, M. T., Chowdhury, A., Sottini, A., Nüesch, M. F., Nettels, D., Best, R. B., & Schuler, B. Ultrafast molecular dynamics observed within a dense protein condensate. bioRxiv.12.12.520135 (2022).

43. Xue, B., Dunbrack, R. L., Williams, R. W., Dunker, A. K., & Uversky, V. N. PONDR-FIT: A meta-predictor of intrinsically disordered amino acids. Biochim. Biophys. Acta 1804, 996–1010 (2010).

44. Uversky, V. N., Gillespie, J. R., & Fink, A. L. Why are “natively unfolded” proteins unstructured under physiologic conditions? Proteins 15, 415–27 (2000).

45. Holehouse, A. S., Das, R. K., Ahad, J. N., Richardson, M. O., & Pappu, R. V. CIDER: Resources to Analyze Sequence-Ensemble Relationships of Intrinsically Disordered Proteins. Biophys. J. 10, 16–21 (2017).

46. Newcombe, E. A., Ruff, K. M., Sethi, A., Ormsby, A. R., Ramdzan, Y. M., Fox, A., Purcell, A. W., Gooley, P. R., Pappu, R. V., & Hatters, D. M. Tadpole-like Conformations of Huntingtin Exon 1 Are Characterized by Conformational Heterogeneity that Persists regardless of Polyglutamine Length. J. Mol. Biol. 11, 1442–1458 (2018).

47. Perdikari, T. M., Jovic, N., Dignon, G. L., Kim, Y. C., Fawzi, N. L., & Mittal, J. A predictive coarse-grained model for position-specific effects of post-translational modifications. Biophys. J. 6, 1187–1197 (2021).

48. Müller, B. K., Zaychikov, E., Bräuchle, C., & Lamb, D. C. Pulsed interleaved excitation. Biophys. J. 89, 3508–22 (2005).

49. Majumdar, A., Mukhopadhyay, S. Fluorescence Depolarization Kinetics to Study the Conformational Preference, Structural Plasticity, Binding, and Assembly of Intrinsically Disordered Proteins. Methods Enzymol. 611, 347–381 (2018).

50. Rai, S. K., Khanna, R., Avni, A., & Mukhopadhyay S. Heterotypic electrostatic interactions control complex phase separation of tau and prion into multiphasic condensates and co-aggregates. Proc. Natl. Acad. Sci. USA 120, (2023).

51. Dogra, P., Joshi, A., Majumdar, A., & Mukhopadhyay, S. Intermolecular Charge-Transfer Modulates Liquid-Liquid Phase Separation and Liquid-to-Solid Maturation of an Intrinsically Disordered pH-Responsive Domain. J. Am. Chem. Soc. 141, 20380–20389 (2019).

52. Ray, S., Singh, N., Kumar, R., Patel, K., Pandey, S., Datta, D., Mahato, J., Panigrahi, R., Navalkar, A., Mehra, S., Gadhe, L., Chatterjee, D., Sawner, A. S., Maiti, S., Bhatia, S., Gerez, J. A., Chowdhury, A., Kumar, A., Padinhateeri, R., Riek, R., Krishnamoorthy, G., & Maji, S. K. α-Synuclein aggregation nucleates through liquid-liquid phase separation. Nat. Chem. 12, 705–716 (2020).

53. Patel, A., Lee, H. O., Jawerth, L., Maharana, S., Jahnel, M., Hein, M. Y., Stoynov, S., Mahamid, J., Saha, S., Franzmann, T. M., Pozniakovski, A., Poser, I., Maghelli, N., Royer, L. A., Weigert, M., Myers, E. W., Grill, S., Drechsel, D., Hyman, A. A., & Alberti, S. A Liquid-to-Solid Phase Transition of the ALS Protein FUS Accelerated by Disease Mutation. Cell 27, 1066–77 (2015).

54. Rhine, K., Makurath, M. A., Liu, J., Skanchy, S., Lopez, C., Catalan, K. F., Ma, Y., Fare, C. M., Shorter, J., Ha, T., Chemla, Y. R., & Myong, S. ALS/FTLD-Linked Mutations in FUS Glycine Residues Cause Accelerated Gelation and Reduced Interactions with Wild-Type FUS. Mol. Cell 19, 666–681(2020).

55. Bolognesi, B., Lorenzo Gotor, N., Dhar, R., Cirillo, D., Baldrighi, M., Tartaglia, G. G., & Lehner, B. A Concentration-Dependent Liquid Phase Separation Can Cause Toxicity upon Increased Protein Expression. Cell Rep. 16, 222–231(2016).

56. Agarwal, A., Rai, S. K., Avni, A., & Mukhopadhyay, S. An intrinsically disordered pathological prion variant Y145Stop converts into self-seeding amyloids via liquid-liquid phase separation. Proc. Natl. Acad. Sci. USA 9, (2021).

57. Avni, A., Joshi, A., Walimbe, A., Pattanashetty, S. G., & Mukhopadhyay, S. Single-droplet surface-enhanced Raman scattering decodes the molecular determinants of liquid-liquid phase separation. Nat. Commun. 28, (2022).

58. Rygula, A., Majzner, K., Marzec, K. M., Kaczor, A., Pilarczyk, M., & Baranska, M. Raman spectroscopy of proteins: a review. J. Raman Spectrosc. 44, 1061–1076 (2013).

59. Tuma, R. Raman spectroscopy of proteins: from peptides to large assemblies. J. Raman Spectrosc. 36, 307–319 (2005).

60. Farag, M., Cohen, S. R., Borcherds, W. M., Bremer, A., Mittag, T., & Pappu, R.V. Condensates formed by prion-like low-complexity domains have small-world network structures and interfaces defined by expanded conformations. Nat. Commun. 13, (2022).

61. Zheng, W., Dignon, G. L., Jovic, N., Xu, X., Regy, R. M., Fawzi, N. L., Kim, Y. C., Best, R. B., & Mittal, J. Molecular Details of Protein Condensates Probed by Microsecond Long Atomistic Simulations. J. Phys. Chem. B. 24, 11671–11679 (2020).

62. Melo, A. M., Coraor, J., Alpha-Cobb, G., Elbaum-Garfinkle, S., Nath, A., & Rhoades, E. A. Functional role for intrinsic disorder in the tau-tubulin complex. Proc. Natl. Acad. Sci. USA 13, 14336–14341 (2016).

63. Wen, J., Hong, L., Krainer, G., Yao, Q. Q., Knowles, T. P. J., Wu, S., & Perrett, S. Conformational Expansion of Tau in Condensates Promotes Irreversible Aggregation. J. Am. Chem. Soc. 25, 13056–13064 (2021).

64. Weickert, S., Wawrzyniuk, M., John, L. H., Rüdiger, S. G. D., & Drescher, M. The mechanism of Hsp90-induced oligomerizaton of Tau. Sci. Adv. 6, (2020).

