## Supplementary Information for "Single-Molecule FRET Illuminates Structural Subpopulations and Dissects Crucial Molecular Events During Phase Separation of a Prion-Like Low-Complexity Domain"

**Table of Contents** **Page number**

1. Materials........................................................ S2
2. Methods........................................................ S2
3. Supplementary Tables........................................................ S10
4. Supplementary Figures........................................................ S13
5. Supplementary References........................................................ S18

#### Materials

Sodium phosphate monobasic dihydrate, sodium phosphate dibasic dihydrate, 2-mercaptoethanol (BME), 1,4-dithiothreitol (DTT), Tris(2-carboxyethyl)phosphine hydrochloride (TCEP), and Urea were of MB grade purity, procured from Sigma (St. Louis, MO, USA). Luria Bertani Broth, Miller (LB), N-cyclohexyl-3-aminopropanesulfonic acid (CAPS), sodium chloride, ethylenediaminetetraacetic acid (EDTA), and nickel chloride were procured from HiMedia Laboratories (MB grade). Kanamycin and isopropyl-β-thiogalactopyranoside (IPTG) were obtained from Gold Biocom (USA). Fluorescent probes like fluorescein-5-maleimide (F-5-M), AlexaFluor488 succinimidyl ester, AlexaFluor488, and AlexaFluor594-maleimide were purchased from Molecular Probes, Invitrogen. Ni-NTA resin was purchased from Qiagen. Amicon membrane filters for concentrating protein were obtained from Merck Millipore. PD-10, NAP-10, and HiLoad 16/600 Superdex-G-200 columns were purchased from GE Healthcare Life Sciences (USA). High-purity milli-Q water was used to prepare all the buffers in this study. A Metrohm 827 lab pH meter was used to adjust the pH (± 0.01) of all the buffer solutions prepared at 25 °C, and all the buffer solutions were filtered before use.

#### Methods

##### Bioinformatics analyses

Various bioinformatics tools were used for the sequence characterization of FUS-LC. Classification of Intrinsically Disordered Ensemble Regions (CIDER) (https://pappulab.wustl.edu/CIDERinfo.html)[1] for visualization of charged and hydrophobic amino acids and Predictor of Natural Disordered Regions (PONDR) (http://www.pondr.com/)[2] for disorder propensity prediction were used. To determine the phase separation propensity catGranule (http://www.tartaglialab.com/)[3] and PScore (http://abragam.med.utoronto.ca/~JFKlab/)[4] (based on π-π interaction propensity) were used. Disorder and phase separation propensity plots were generated using the Origin software.

##### Construct details and Site-directed mutagenesis

All single cysteine and disease mutants were created by site-directed mutagenesis using the recombinant MBP-$His_6$-FUS-LC WT (Addgene plasmid # 98653; https://www.addgene.org/98653/; RRID: Addgene_98653) cloned in pTHMT vector which was a kind gift from Nicolas L. Fawzi. The primer sets used for introducing these point mutations have been listed in Supplementary Table 1. All mutations were confirmed by sequencing.

**Recombinant protein expression and purification**

Wild-type and all the variants of FUS-LC were transformed in *E. coli* BL21(DE3) std cells, overexpressed, and purified using affinity chromatography, followed by gel-filtration chromatography. Bacterial cultures were grown at 37 °C, 220 rpm, to an $O.D._{600}$ of 0.8-1. Protein overexpression was induced by adding 1 mM isopropyl-β-thiogalactopyranoside (IPTG) and further growing cultures at 37 °C for 4-5 h. Bacterial cells were harvested by centrifugation at 4 °C, 3220 x g for 30 minutes, and stored at -80 °C for future use. Cell pellets were resuspended in lysis buffer (20 mM sodium phosphate, 300 mM NaCl, 10 mM imidazole, pH 7.4) and were lysed by probe sonication at 5% amplitude, 15 seconds ON, and 10 seconds OFF for 20 minutes. The lysate was centrifuged at 4 °C, 15,557 x g for 1 h to remove the cell debris, and the supernatant was loaded onto a Ni-NTA column. The column was washed, and the bound protein was eluted with elution buffer (20 mM sodium phosphate, 300 mM NaCl, 300 mM imidazole, pH 7.4).

The N-terminal MBP-$His_6$ tag was cleaved by adding recombinantly expressed and in-house purified TEV protease at a 1:40 molar ratio (TEV: protein), followed by incubation at 30 °C for 1.5 h. It was then subjected to overnight dialysis at room temperature. The cleaved protein was passed through Ni-NTA column to separate the uncleaved species and TEV protease and flowthrough were collected and concentrated using a 10 kDa MWCO Amicon filter. Concentrated protein was further loaded on HiLoad 16/600 Superdex-G-200 (GE) column equilibrated with the SEC buffer (20 mM CAPS, 150 mM NaCl, pH 11). SEC elution fractions were run on an SDS-PAGE gel to determine fractions containing protein of interest. Pure protein fractions were pooled, concentrated, and buffer-exchanged into 20 mM CAPS, pH 11 buffer using a PD-10 column. Pure protein was concentrated using a 3 kDa MWCO Amicon filter, and concentration was estimated by measuring absorbance at 280 nm ($\varepsilon_{280}$ = 30,720). Pure protein was flash-frozen and stored at - 80 °C.

**Circular dichroism (CD) measurements**

Far-UV CD spectra were recorded using a Chirascan spectrophotometer (Applied Photophysics, UK) in a quartz cuvette of 1 mm path length. Wild-type and G156E FUS-LC were diluted to 10 μM in 20 mM sodium phosphate buffer, pH 7.4. Measurements were made for buffer and monomeric FUS-LC. Recorded absorption spectra were averaged over 10 scans, followed by blank subtraction using ProData software, and plotted using the Origin software.

**Phase separation assays**

Protein stock was thawed on ice and diluted up to 200 µM in 20 mM phosphate buffer, pH 7.4. Phase separation of wild-type and G156E FUS-LC was induced by the addition of 250 mM NaCl in the reaction mixture. Spontaneous phase separation of FUS-LC into liquid droplets was indicated by the immediate rise in turbidity upon mixing with salt.

**Turbidity assay**

The turbidity of monomeric FUS-LC and phase-separated samples of wild-type and G156E FUS-LC were monitored by measuring absorbance at 350 nm on a Multiskan Go (Thermo Scientific) plate reader. Droplet reactions of 100 µL (200 µM FUS-LC in 20 mM phosphate, 250 mM NaCl, pH 7.4) were set up and used for the turbidity measurements. The mean and standard errors were obtained from at least 3 independent sets of measurements.

**Fluorescence labeling**

Single-cysteine FUS-LC variants were labeled with fluorescein-5-maleimide (F-5-M) and AlexaFluor488-C5-maleimide under denaturing buffer conditions (8 M Urea, 20 mM phosphate, pH 7.5) for anisotropy and FRAP measurements. Pure protein was incubated with 0.3 mM tris(2-carboxyethyl)phosphine (TCEP) for 30 minutes on ice and was mixed with fluorescent dyes in a molar ratio of 1:30 (for F-5-M) and 1:3 (for AlexaFluor488-maleimide). The labeling mixture was incubated in the dark under stirring conditions at room temperature for 3 h. Following the reaction, the excess free dye was removed by buffer exchange using a NAP-10 column. For dual-labeling of FUS-LC single-cysteine variants, the pure protein was incubated under denaturing conditions (8 M Urea, 20 mM phosphate, pH 8) with the amine-reactive NHS ester of AlexaFluor488 (donor dye) in a molar ratio of 1:4 under shaking at 25 °C for 4 h. The unreacted dye was further removed using a NAP-10 column, and the eluted protein was concentrated and used for labeling with the thiol-reactive acceptor dye. The donor-labeled protein was mixed with AlexaFluor594-maleimide in a ratio of 1:4 and incubated at 25 °C with stirring for 5 h, under denaturing conditions (8 M Urea, 20 mM phosphate, pH 7.5). The labeling reaction was then buffer exchanged with a NAP-10 column, and the remaining free dye was removed using a 3 kDa MWCO Amicon filter. All the single and dual-labeled proteins were concentrated using a 3 kDa MWCO Amicon filter. Labeling efficiencies were estimated by measuring absorbance at 280 nm ($\varepsilon_{280nm}$ = 30,720 $M^{-1}cm^{-1}$, for FUS-LC cysteine variants), 494 nm ($\varepsilon_{494}$ = 73,000 $M^{-1}$ $cm^{-1}$, for AlexaFluor488 and $\varepsilon_{494}$ = 68,000 $M^{-1}$ $cm^{-1}$, for F-5-M) and 590 nm ($\varepsilon_{590}$ = 92,000 $M^{-1}$ $cm^{-1}$ for AlexaFluor594) to estimate the total protein and labeled protein concentrations.

**Confocal microscopy**

All fluorescence microscopy imaging was performed on ZEISS LSM 980 Elyra 7 super-resolution Microscope using a 63x oil-immersion objective (N.A. 1.4) and a monochrome cooled high-resolution AxioCamMRm Rev. 3 FireWire(D) camera. Phase separation of 200 μM unlabeled FUS-LC was induced in the presence of 0.1% AlexaFluor488-labeled FUS-LC by the addition of 250 mM NaCl (20 mM phosphate, pH 7.4). Reactions were incubated at room temperature for 5 minutes and a 5-10 μL sample was placed on a glass coverslip and imaged using a 488 nm laser diode (11.9 mW). For two-color imaging, the droplet reaction was spiked with 0.05% of dual-labeled FUS-LC and imaged using 488 nm and 590 nm excitation sources, respectively. The images were acquired at 1840 x 1840 pixels and 16-bit depth resolution. Airyscan images of the fluorescently labeled droplets were acquired by utilizing the confocal laser scanning microscope via the Airyscan 2 detector equipped with 32 channels (GaAsP). Image processing and analyses were performed on in-built instrument software Zen Blue 3.2 and ImageJ (NIH, Bethesda, USA).

**Fluorescence recovery after photobleaching (FRAP) measurements**

FRAP experiments were done on ZEISS LSM 980 Elyra 7 super-resolution microscope using a 63x oil-immersion objective (N.A 1.4) and a monochrome cooled high-resolution AxioCamMRm Rev. 3 FireWire(D) camera. A region of 1 μm was bleached inside droplets doped with 0.1% AlexaFluor488-labeled FUS-LC using a 488 nm laser diode. The recovery was recorded using the Zen Blue 3.2 (ZEISS) software. FRAP measurements were performed for at least 8 independent droplets for both wild-type and G156E FUS-LC. Fluorescence recovery curves were normalized, background corrected, and plotted using the Origin software.

**Steady-state fluorescence measurements**

Steady-state FRET experiments were performed on a Fluoromax-4 spectrofluorometer (Horiba Jobin Yvon, NJ, USA) using a 1-mm pathlength quartz cuvette. For all the experiments, 50 nM of dual-labeled FUS-LC variants were used. The donor fluorophore (AlexaFluor488) was excited at 494 nm and fluorescence emission was recorded from 515 nm to 700 nm to monitor both donor and acceptor emission spectra.

**Single-droplet FRET imaging by acceptor photobleaching**

Phase separation of 200 μM FUS-LC (20 mM phosphate, pH 7.4) was induced by the addition of 250 mM NaCl in salt in the presence of 0.05 % dual-labeled single-cysteine variants of FUS-LC. The droplet reaction was imaged on a ZEISS LSM 980 Elyra 7 super-resolution microscope using a 63x oil-immersion objective (N.A 1.4) and a monochrome-cooled high-resolution AxioCamMRm Rev. 3 FireWire(D) camera. To determine the FRET efficiency, the acceptor present in a dual-labeled droplet was photobleached using a 594 nm laser, and the increase in the donor fluorescence intensity was recorded upon bleaching the acceptor fluorophore. FRET efficiencies were estimated using the Zen Blue 3.2 (ZEISS) software.

**Dynamic light scattering (DLS)**

For estimating the hydrodynamic radii of monomeric FUS-LC, a dynamic light scattering instrument (Malvern Zetasizer) was used. All the reaction buffers were filtered using 0.02 µm filters. Monomeric FUS-LC (50 µM in 20 mM phosphate, pH 7.4) in the absence and presence of 250 mM NaCl was used for measurements at room temperature.

**$C_{sat}$ estimation**

Droplet reactions (200 µM FUS-LC) were induced by the addition of 250 mM NaCl ( in 20 mM sodium phosphate, pH 7.4 buffer) and incubated at 25 °C for 10 minutes. The reactions were then subjected to ultracentrifugation at 25 °C, 18000 x g for 30 minutes. The supernatant was removed carefully without disturbing the pellet to estimate the dilute phase concentration. The protein saturation concentration ($C_{sat}$) of the dilute phase was estimated by measuring the absorbance at 280 nm. ($\varepsilon_{280}$ = 30,720)

**Single-molecule FRET experiments and data analysis**

In single-molecule FRET experiments, the ratiometric FRET efficiency (E) for each molecule is recorded from the fluorescence bursts that are separated into donor ($I_D$) and acceptor ($I_A$) signals using the following equation.

$$E = \frac{1}{1+\left(\frac{I_D}{I_A}\right)\gamma} \quad \text{--- (1)}$$

where $\gamma$ is a correction factor obtained from different quantum yields of donor and acceptor dyes and the detection efficiencies for donor and acceptor channels.

Single-molecule FRET experiments were performed using MicroTime 200 time-resolved confocal microscope (PicoQuant) in a pulsed interleaved excitation (PIE) mode. All the single-molecule FRET experiments were performed in 20 mM phosphate buffer, 250 mM NaCl, pH 7.4. Measurements in the monomeric dispersed phase were performed in the presence of 75-150 pM of dual-labeled FUS-LC, and droplet formation of 200 µM FUS-LC was performed in the presence of 5-10 pM of dual-labeled protein. Reactions were set in a buffer containing n-propyl gallate as an oxygen scavenger to improve the photostability of the fluorophore in the solution[5]. Pulsed laser sources (485 nm and 594 nm) at a frequency of 20 MHz were used to alternately excite the donor and the acceptor fluorophores within the dual-labeled samples using a 60x water-immersion objective (N.A.= 1.2). The laser power was fixed at 45-60 μW (40-50 μW for 485 nm and 5-10 μW for 594 nm laser) measured at the back aperture of the objective for the dispersed phase and 5.5-8.5 μW (5-7 μW for 485 nm and 0.5-1.5 μW for 594 nm laser), in order to minimize the saturation, background counts, and photobleaching of the acceptor. The lasers were focused inside the solution (50 μm from the surface) for the dispersed phase and within single droplets (2-4 μm inside) for the condensed phase to obtain fluorescence emission bursts. The emitted photons were collected and focused through a 50 μm pinhole, and a dichroic beam splitter (zt594rdc) was used to separate the donor and acceptor emission. The emission was filtered (BP 535/50 nm for the green channel and LP 594 for the red channel) and detected by the respective single-photon avalanche diode (SPAD) detectors. Data were collected and analyzed using the SymphoTime64 software v2.7. A typical binning time used was 0.5 ms and 1 ms, and using PIE, the bursts containing both donor and acceptor signals were considered for FRET analysis. The donor and acceptor counts were corrected for the background with a minimum threshold of 35 photons used for the further selection of bursts to construct the FRET efficiency histogram. The FRET efficiencies were corrected for the spectral crosstalk between the donor and acceptor fluorophore ($\alpha = 0.05$), direct excitation of the acceptor by donor laser ($\beta = 0.003$), and difference in the detection efficiencies of the donor and acceptor channels ($\gamma = 1.12$), estimated as described previously[6,7]. All the FRET efficiency histograms were constructed for > 20,000 events. The FRET efficiency histograms were plotted and fitted using a Gaussian peak function in the Origin software.

**Fluorescence correlation spectroscopy (FCS)**

FCS measurements were performed using the same MicroTime 200 setup for the monomeric phase and the condensed phase of wild-type and G156E FUS-LC. A free dye solution of 1 nM AlexaFluor488 was used to estimate the structure parameter of confocal volume (5.52), which was used for further FCS analyses. For the dispersed phase measurements, data were acquired in the presence of 10 nM AlexaFluor488-labeled FUS-LC in the presence of salt (250 mM NaCl). For single-droplet FCS measurements, droplet reactions were set up with 1-3 nM AlexaFluor488 labeled FUS-LC, and these samples were placed on glass coverslips. Measurements were performed by focusing inside the solution for the dispersed phase and within single droplets for the condensed phase of wild-type and G156E FUS-LC. FCS data were collected, analyzed, and fitted with the triplet-state model using SymphoTime64 software v2.7, as previously described by us[8].

**Single-droplet steady-state and time-resolved fluorescence anisotropy measurements**

MicroTime 200 time-resolved confocal microscope (PicoQuant) was used for performing steady-state fluorescence anisotropy measurements of dispersed and droplet phase of FUS-LC spiked, 0.1% F-5-M-labeled protein with single-cysteine variants at residue positions 16, 86, 108, and 148. Fluorescein-5-maleimide (F-5-M) dye was used as a thiol-reactive anisotropy probe owing to its short linker length which accurately reports on the rotational dynamics of the polypeptide chain. Freshly phase-separated droplet reactions were spotted on a coverslip with a thickness of 1.5 mm placed directly on a Super Apochromat 60x water immersion objective with 1.2 NA (Olympus). Samples were excited with the 485 nm laser, and the emitted fluorescence was collected and filtered by a bandpass emission filter (BP 535/50) before entering the pinhole (50 μm). The in-focus emitted light exiting the pinhole was split into the two detector channels by a polarizing beam-splitter placed before the detectors and detected by the respective Single-Photon Avalanche Diodes (SPADs). Anisotropy imaging was performed for single-droplet steady-state anisotropy measurements, and a point time trace was obtained for time-resolved anisotropy measurements. The correction factors were calculated by performing fluorescence measurements in a free dye solution and utilized to estimate steady-state anisotropy using the commercially available SymphoTime64 software v2.7. The fluorescence anisotropy ($r_{ss}$) is given by

$$r_{ss} = \frac{I_{\parallel} - I_{\perp}}{[1-3\mathrm{L2}]I_{\parallel} + [2-3\mathrm{L1}]I_{\perp}} \text{ --- (2)}$$

where $I_{\parallel}$ and $I_{\perp}$ are the background corrected parallel and perpendicular fluorescence intensities, and L1 (0.308) and L2 (0.0368) are the objective correction factors[9].

For time-resolved fluorescence anisotropy decay analysis, the decay profiles obtained from SymphoTime64 software v2.7 was further analyzed by global fitting using the following relationships.

$$I_{\parallel}(t) = 1/3I(t)[1 + 2r(t)] \text{ --- (3)}$$

$$I_{\perp}(t) = 1/3I(t)[1 - r(t)] \text{ --- (3)}$$

where $I_{\parallel}(t)$, $I_{\perp}(t)$, and I(t) denote the time-dependent fluorescence intensities collected at the parallel, perpendicular, and magic angle (54.7°) geometry. The perpendicular component was always corrected using the G-factor that was intendedly obtained from free dye in the buffer. The time-resolved fluorescence anisotropy decay profiles were fitted using a biexponential decay model yielding two rotational correlation times, namely, fast ($\phi_1$) and slow ($\phi_2$) rotational correlation times as follows.

$$r(t) = r_0[\beta_1 e^{\left(\frac{-t}{\phi_1}\right)} + \beta_2 e^{\left(\frac{-t}{\phi_2}\right)}] \text{ --- (4)}$$

where $r_0$ denotes the (time-zero) fundamental anisotropy of the fluorophore, and $\beta_1$ and $\beta_2$ the amplitudes associated with $\phi_1$ and $\phi_2$, respectively. The goodness of fit was estimated based on the autocorrelation function, randomness of residuals, and reduced $\chi^2$ values[10].

**Raman spectroscopy**

Raman spectra of phase-separated individual droplets of wild-type and G156E FUS-LC were acquired on an inVia laser Raman microscope (Renishaw, UK) at room temperature. Freshly phase-separated samples (3-5 μL) were drop cast onto a glass slide covered with aluminum foil. Single droplets were focused using a 100x long working distance objective lens (Nikon, Japan). The samples were excited with an NIR laser (785 nm) with an exposure time of 10 seconds at a laser power of 500 mW (100%), and an edge filter of 785 nm was used to block the Rayleigh scattering. The collected Raman scattering was dispersed using a diffraction grating (1200 lines/mm) and further detected by an air-cooled CCD detector. Data were acquired for 5 accumulations, after which collected spectra were background corrected and smoothened using inbuilt software Wire 3.4. All the data were plotted and analyzed using the Origin software.

**Supplementary Table 1.** Primers used for site-directed mutagenesis.

| | |
|---|---|
| A16C Forward | CCCAAAGCTATGGGTGCTACCCCACCCAGC |
| A16C Reverse | GCTGGGTGGGGTAGCACCCATAGCTTTGGG |
| S86C Forward | CTATGGCAGTAGCCAGTGCTCCCAATCGTC |
| S86C Reverse | GACGATTGGGAGCACTGGCTACTGCCATAG |
| S108C Forward | CCAGCTCCCAGCTGCACCTCGGGAA |
| S108C Reverse | TTCCCGAGGTGCAGCTGGGAGCTGG |
| S148C Forward | AAAGCTATGGACAGCAGCAATGCTATAATCCCCC |
| S148C Reverse | GGGGGATTATAGCATTGCTGCTGTCCATAGCTTT |
| LC G156E Forward | GCTATAATCCCCCTCAGGGCTATGAACAGCAGAACCAGTACAACAGC |
| LC G156E Reverse | GCTGTTGTACTGGTTCTGCTGTTCATAGCCCTGAGGGGGATTATAGC |

**Supplementary Table 2.** Observed FRET efficiencies varying the inter-residue length estimated from single-molecule FRET data analyses and the comparison with the calculated FRET efficiencies based on the random coil model[11].

| Constructs | Number of residues | Experimental FRET efficiencies from single-molecule FRET studies | Calculated FRET efficiencies |
|---|---|---|---|
| N-to-86 | 86 | 0.76 ± 0.02 | 0.32 |
| N-to-108 | 108 | Subpopulation 1: 0.80 ± 0.02<br>Subpopulation 2: 0.97 ± 0.03 | 0.16 |
| N-to-148 | 148 | Subpopulation 1: 0.73 ± 0.01<br>Subpopulation 2: 0.09 ± 0.02 | 0.06 |

**Supplementary Table 3.** FRET efficiencies varying the inter-residue length estimated from single-molecule FRET data analyses for wild-type and G156E FUS-LC condensates.

| Constructs | FRET efficiencies from single-droplet single-molecule FRET studies for wild-type FUS-LC | FRET efficiencies from single-droplet single-molecule FRET studies for G156E FUS-LC |
|---|---|---|
| N-to-86 | 0.64 ± 0.01 | 0.64 ± 0.01 |
| N-to-108 | Subpopulation 1: 0.79 ± 0.08<br>Subpopulation 2: 1.00 ± 0.01 | Subpopulation 1: 0.86 ± 0.04<br>Subpopulation 2: 1.01 ± 0.01 |
| N-to-148 | Subpopulation 1: 0.30 ± 0.01<br>Subpopulation 2: 0.90 ± 0.02 | Subpopulation 1: 0.24 ± 0.02<br>Subpopulation 1: 0.79 ± 0.06 |

**Supplementary Table 4.** Rotational correlation times and associated amplitudes recovered by fitting fluorescence anisotropy decay kinetics using a biexponential decay model for the monomeric dispersed phase and individual droplets.

| **Residue position** | | **Fast rotational correlation time ($\phi_1$) and amplitude ($\beta_1$)** | **Slow rotational correlation time ($\phi 2$) and amplitude ($\beta 2$)** |
|---|---|---|---|
| **16** | Monomer | 0.95 ± 0.12 ns (0.66 ± 0.08) | 4.34 ± 0.45 ns (0.33 ± 0.08) |
| | Droplet | 1.02 ± 0.16 ns (0.29 ± 0.12) | 57.73 ± 5.44 ns (0.74 ± 0.01) |
| **86** | Monomer | 0.77 ± 0.07 ns (0.66 ± 0.05) | 4.11 ± 0.32 ns (0.33 ± 0.05) |
| | Droplet | 1.42 ± 0.09 ns (0.26 ± 0.00) | 62.49 ± 4.29 ns (0.73 ± 0.00) |
| **108** | Monomer | 0.93 ± 0.06 ns (0.65 ± 0.02) | 5.61 ± 0.81 ns (0.34 ± 0.02) |
| | Droplet | 1.11 ± 0.08 ns (0.23 ± 0.01) | 58.26 ± 1.37 ns (0.76 ± 0.01) |
| **148** | Monomer | 0.92 ± 0.06 ns (0.79 ± 0.05) | 5.16 ± 0.70 ns (0.20 ± 0.05) |
| | Droplet | 1.29 ± 0.24 ns (0.21 ± 0.02) | 63.50 ± 3.92 ns (0.78 ± 0.02) |

### Supplementary Figures

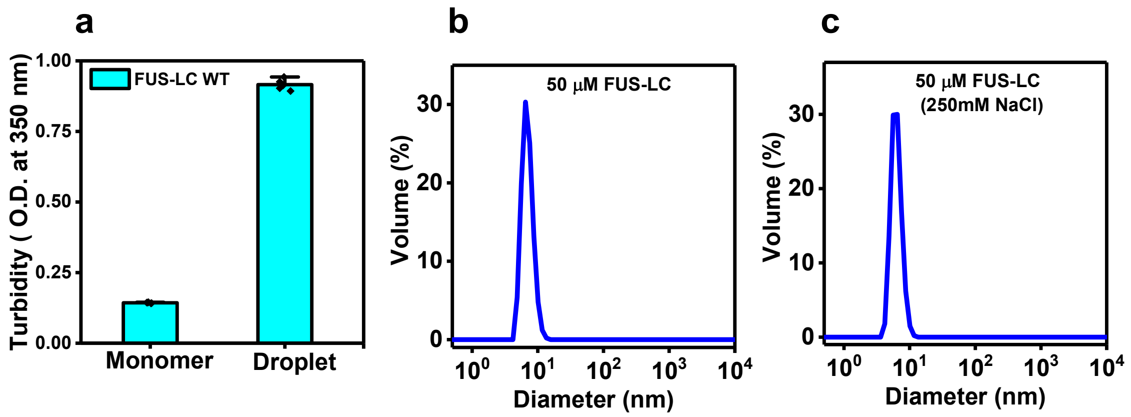


**Supplementary Figure 1**. a. Turbidity plot of wild-type FUS-LC measured at 350 nm in monomeric and droplet conditions. Turbidity measurements were performed for 200 μM of FUS-LC in 20 mM phosphate buffer, pH 7.4 without salt and with 250 mM NaCl for monomer and droplet phases, respectively. Data represent mean ± SD (n = 5). Distribution of particle size obtained from dynamic light scattering (DLS) measurements of monomeric FUS-LC (50 μM protein, in 20 mM phosphate, pH 7.4) (n = 3) under non-phase separating (no salt) (b) and phase-separating conditions (c) (250 mM NaCl).

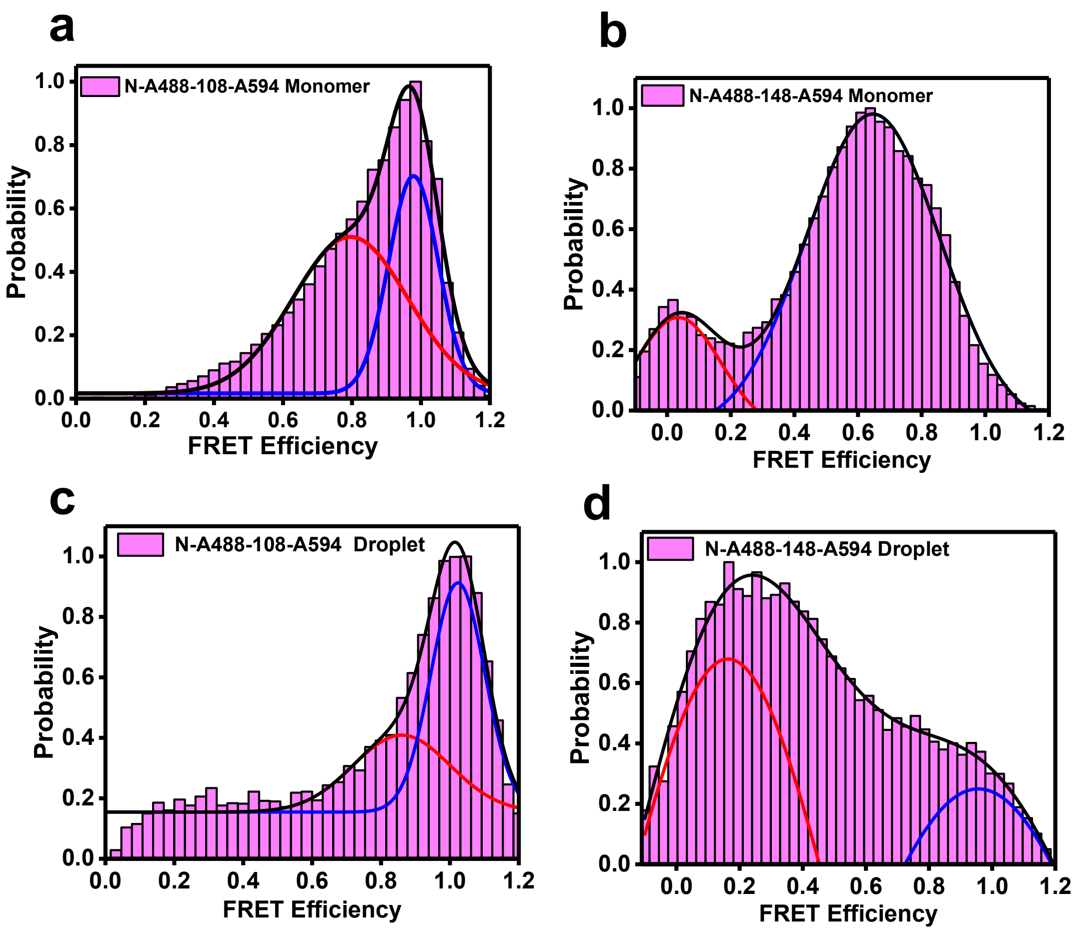


**Supplementary Figure 2.** The effect of binning time showing no significant changes in the FRET efficiency histograms. These results with a 1 ms binding time are similar to the results with 0.5 ms binning time shown in Figure 3. Single-molecule FRET efficiency histograms obtained for the dispersed phase of N-to-108 (a) and N-to-148 (b) constructs acquired in the presence of 75-150 pM dual-labeled FUS-LC. Single-droplet single-molecule FRET efficiency histograms obtained for the condensed phase of N-to-108 (c) and N-to-148 (d) constructs acquired in the presence of 5-10 pM dual-labeled FUS-LC. All the single-molecule FRET measurements were performed in 20 mM phosphate buffer, 250 mM NaCl, pH 7.4, the donor and acceptor emissions were recorded in PIE mode and FRET efficiency distribution was obtained with a binning time of 1 ms using SymphoTime64 software v2.7.

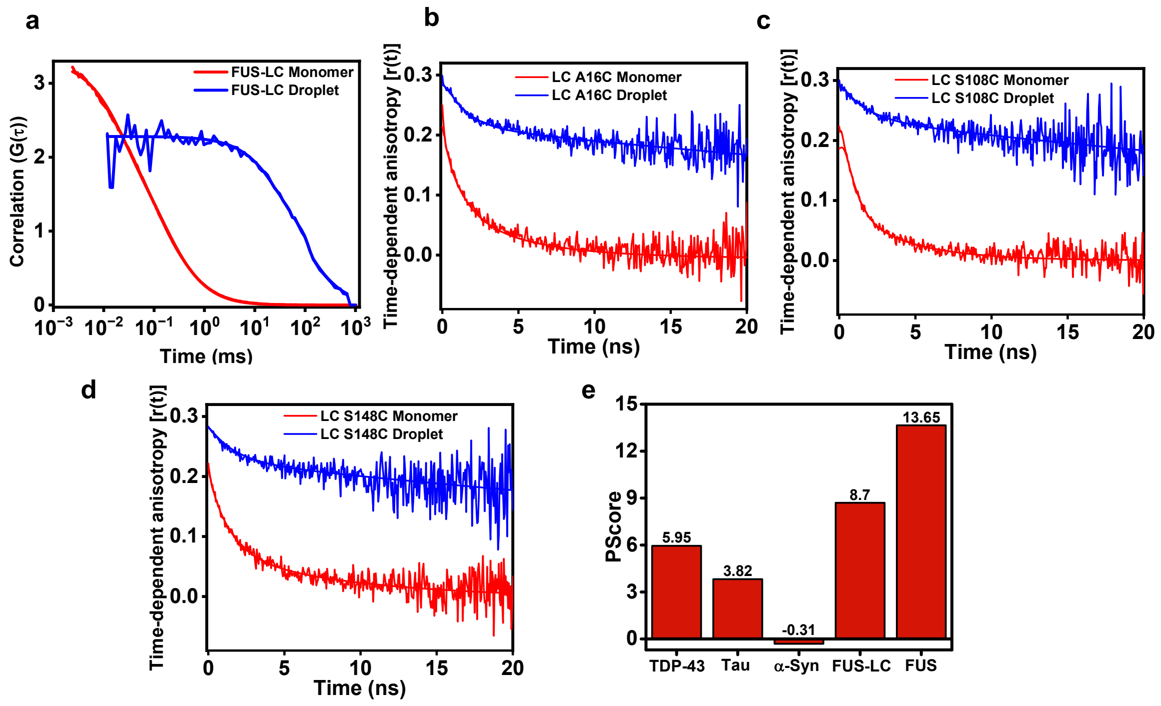


**Supplementary Figure 3.** a. Unnormalized FCS autocorrelation plots with fits for the monomeric and condensed phase of wild-type FUS-LC in the presence of 10 nM (monomer) and 1-3 nM (droplet) AlexaFluor488-labeled FUS-LC. The normalized FCS plots are shown in Fig. 4a. b-d. Time-resolved fluorescence anisotropy decay of single-cysteine variants of FUS-LC obtained within the dispersed phase and single droplets (F-5-M labeled at residue locations 16, 108, and 148) (b), (c), and (d), respectively. Solid lines are fits obtained from biexponential decay analysis. e. Comparison of Pscore values of FUS-LC with other well-studied phase-separating proteins.

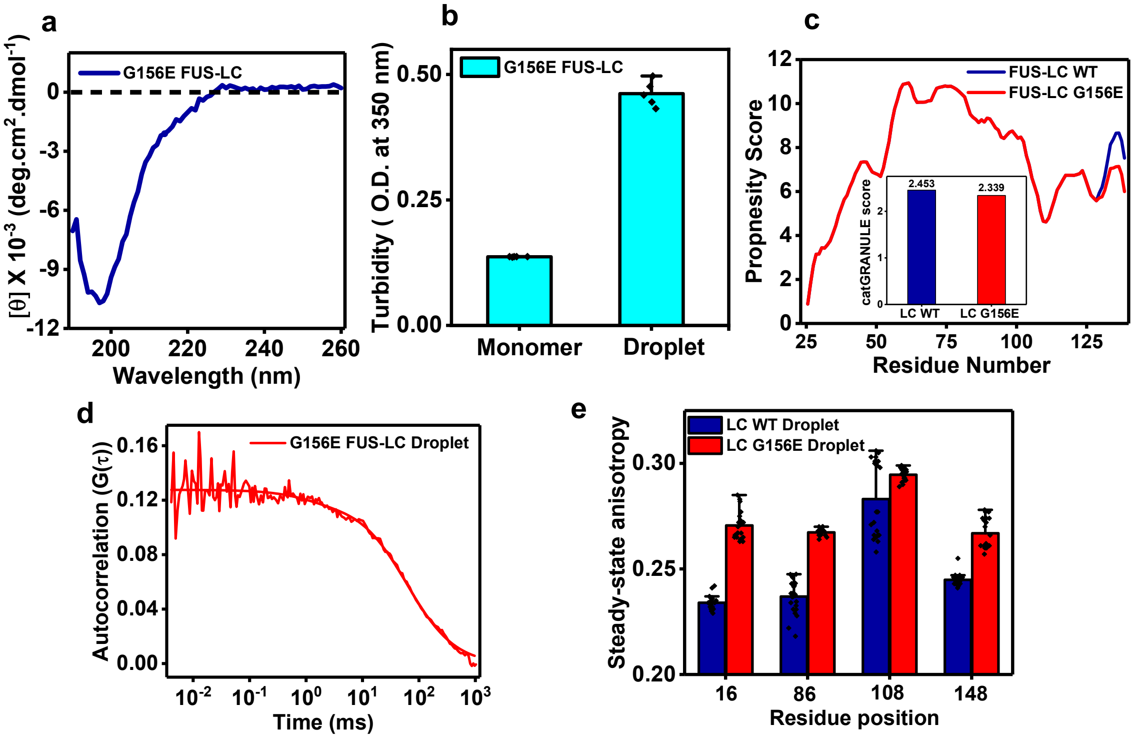


**Supplementary Figure 4.** a. The circular dichroism spectrum of G156E FUS-LC shows similar structural features to wild-type FUS-LC. CD measurements were performed for 10 µM G156E FUS-LC in 20 mM phosphate, pH 7.4. b. The solution turbidity plot of G156E FUS-LC (200 µM protein in 20 mM phosphate, pH 7.4) indicates phase separation in the presence of 250 mM salt. Data represent mean ± SD (n = 5). c. Comparison of phase separation propensity plots of wild-type and mutant G156E FUS-LC using bioinformatics tool catGranule shows a slightly lower propensity of G156E FUS-LC. Inset shows a comparison of catGranule score for wild-type and G156E FUS-LC. d. Autocorrelation plot obtained by FCS measurements within individual condensates of G156E FUS-LC. Droplets of G156E FUS-LC were doped with 1-3 nM AlexaFluor488-labeled FUS-LC for FCS measurements. e. Comparison of single-droplet steady-state fluorescence anisotropy values within wild-type and G156E FUS-LC droplets. Steady-state fluorescence anisotropy at all the positions showed a slight increase within G156E droplets. Data represent mean ± SD for n = 20 (data for wild-type FUS-LC droplets are the same as shown in Fig. 4d and included here for comparison).

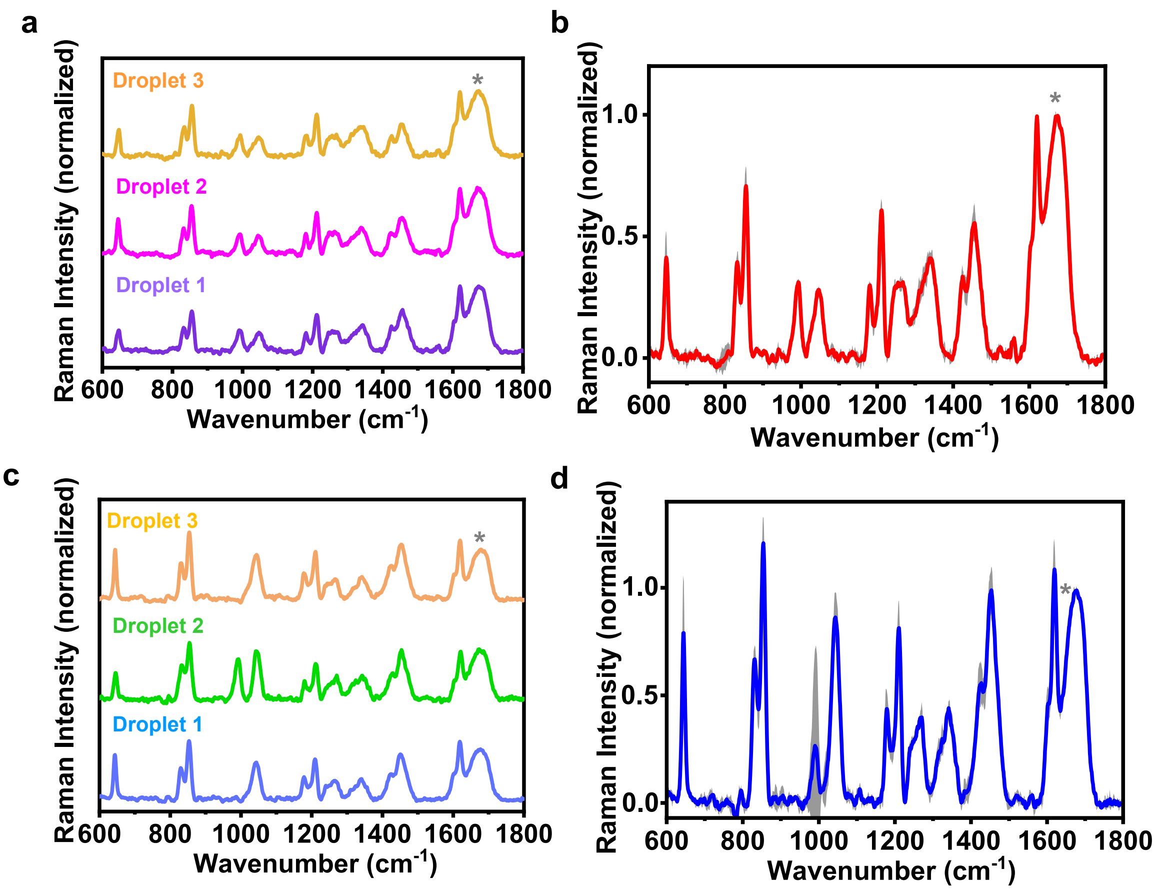


**Supplementary Figure 4.** a. Representative single-droplet normal Raman spectra of individual FUS-LC droplets (spectra recorded at 500 mW laser power, 100x objective; the number of droplets, n = 3). b. Mean and standard deviation of Raman spectra shown in (a). Raman spectra are normalized with respect to the amide I band at ~ 1673 $cm^{-1}$, marked by an asterisk. See "Methods" for details of data acquisition, processing, and analysis. c. Representative single droplet normal Raman spectra of individual G156E FUS-LC droplets (spectra recorded at 500 mW laser power, 100x objective; the number of droplets, n = 3). d. The mean and standard deviation of Raman spectra shown in (c). Raman spectra are normalized with respect to the amide I band at ~ 1673 $cm^{-1}$, marked by an asterisk.

#### References

1. Holehouse, A. S., Das, R. K., Ahad, J. N., Richardson, M. O., & Pappu, R. V. CIDER: Resources to Analyze Sequence-Ensemble Relationships of Intrinsically Disordered Proteins. *Biophys. J.* **10,** 16-21 (2017).
2. Xue, B., Dunbrack, R. L., Williams, R. W., Dunker, A. K., & Uversky, V. N. PONDR-FIT: A meta-predictor of intrinsically disordered amino acids. *Biochim. Biophys. Acta* **1804,** 996–1010 (2010).
3. Bolognesi, B., Lorenzo Gotor, N., Dhar, R., Cirillo, D., Baldrighi, M., Tartaglia, G. G., & Lehner, B. A Concentration-Dependent Liquid Phase Separation Can Cause Toxicity upon Increased Protein Expression. *Cell Rep.* 16, 222–231(2016).
4. Vernon, R. M., Chong, P. A., Tsang, B., Kim, T. H., Bah, A., Farber, P., Lin, H., & Forman-Kay, J. D. Pi-Pi contacts are an overlooked protein feature relevant to phase separation. *eLife* **9**, (2018).
5. Aitken, C. E., Marshall, R. A., & Puglisi, J. D. An oxygen scavenging system for improvement of dye stability in single-molecule fluorescence experiments. *Biophys. J.* **941,** 826-35 (2008).
6. Schuler, B., Lipman, E. A., & Eaton, W. A. Probing the free-energy surface for protein folding with single-molecule fluorescence spectroscopy. *Nature* **419,** 743-7 (2002).
7. Kudryavtsev, V., Sikor, M., Kalinin, S., Mokranjac, D., Seidel, C. A., & Lamb, D. C. Combining MFD and PIE for accurate single-pair Förster resonance energy transfer measurements. *ChemPhysChem* **13,** 1060-78 (2012).
8. Rai, S. K., Khanna, R., Avni, A., & Mukhopadhyay S. Heterotypic electrostatic interactions control complex phase separation of tau and prion into multiphasic condensates and co-aggregates. *Proc. Natl. Acad. Sci. USA.* **120,** (2023).
9. Schaffer, J., Volkmer, A., Eggeling, C., Subramaniam, V., Striker, G., & Seidel, C. A. M. Identification of Single Molecules in Aqueous Solution by Time-Resolved Fluorescence Anisotropy. *J. Phys. Chem. A* **103,** 331–336 (1999).
10. Majumdar, A., Mukhopadhyay, S. Fluorescence Depolarization Kinetics to Study the Conformational Preference, Structural Plasticity, Binding, and Assembly of Intrinsically Disordered Proteins. *Methods Enzymol.* **611,** 347-381 (2018).
11. Melo, A. M., Coraor, J., Alpha-Cobb, G., Elbaum-Garfinkle, S., Nath, A., & Rhoades, E. A. Functional role for intrinsic disorder in the tau-tubulin complex. *Proc. Natl. Acad. Sci. USA* **13,** 14336-14341 (2016).
